# Neural stem cell epigenomes and fate bias are temporally coordinated during corticogenesis

**DOI:** 10.1101/2025.04.03.647013

**Authors:** Yonatan Shapira, Florian Noack, Silvia Vangelisti, Faye Chong, Aviezer Lifshitz, Amos Tanay, Boyan Bonev

**Affiliations:** Department of Molecular Cell Biology, Weizmann Institute of Science, Rehovot, Israel; Department of Computer Science and Applied Mathematics, Weizmann Institute of Science, Rehovot, Israel; Helmholtz Pioneer Campus, Helmholtz Center Munich, Neuherberg, Germany; Physiological Genomics, Biomedical Center, Ludwig-Maximilians-Universität München, Munich, Germany

**Keywords:** Corticogenesis, neural stem cells, single-cell epigenomics, 3D genome, DNA methylation

## Abstract

The cerebral cortex orchestrates complex cognitive functions, yet how its distinct temporal lineages are molecularly patterned during development remains unresolved. Here, we integrate single-cell transcriptomics and chromatin accessibility, together with genome-wide profiling of DNA methylation and 3D chromosomal contact across mouse corticogenesis (E13–E18) to elucidate cell fate transitions. Using metacell flow analysis, we reveal that neural stem cells (NSCs) progressively shift from a progenitor–biased state toward an astrocytic lineage and that this process is accompanied with changes in DNA methylation and 3D genome organization. A model integrating transcription factor motif affinities with epigenetic features identifies key regulators of cis-regulatory element (CRE) activation. *In vivo* reporter assays further decouple the intrinsic regulatory potential of CREs from context-dependent synergistic activation. Collectively, our findings uncover temporal epigenomic reprogramming that underlies the evolving differentiation potential of NSCs, providing insights into the intrinsic and extrinsic mechanisms that pattern cortical lineages.

## Introduction

The cerebral cortex is the region of the brain responsible for cognitive function, sensory perception and consciousness. It contains an unparalleled variety of neuron subtypes, with unique molecular, morphological and connectivity features, which are generated in a precise temporal sequence from neural progenitors^1–5^. Fate-mapping studies have revealed that although the majority of neural stem cells (NSC) are multipotent, a small proportion (10-20%) may be pre-committed to a certain fate early on in development^6,7^. Furthermore, postmitotic refinement of subtype identity has also been observed^8^, which can enhance or counteract lineage initiation in progenitor cells. Single-cell RNA-seq in the mouse^9–11^ and human fetal brain^12–17^ have extensively profiled the remarkable diversity of cell types established during development, but the precise molecular mechanisms involved in lineage specification are still not resolved.

There is increasing evidence that chromatin remodeling and epigenetic regulation are essential for determining fate choices in the cortex^18–20^. Transplantation studies have revealed that the lineage potential of cortical progenitors to generate different types of neurons is primarily cell-autonomous^21,22^ but external stimuli such as bioelectric membrane properties or Wnt signaling can overwrite the inherent developmental programs^22,23^. Additional evidence was provided by the phenotypic defects in the timing and rate of neuronal differentiation in Polycomb mutants^18,20^, disruption of the barrel cortex upon neuron-specific conditional CTCF deletion^24^ and impaired neuronal migration following mutations in the cohesin loading factor Nipbl (associated with Cornelia De Lange syndrome)^25^. Furthermore, the transcription factor (TF) Pax6 has been shown to interact directly with chromatin remodelers^26^, Neurog2 and Ascl1 have been proposed to act as pioneering TFs^27^, while Lhx2 and Ldb1 are essential for enhancer-promoter contacts in olfactory sensory neurons^28^. Finally, global chromatin compaction, possibly mediated by changes in the expression of high mobility group proteins, has been shown to be associated with lineage restriction of neural progenitors during cortical development^29,30^.

We and others have previously shown that cellular identity in neuronal differentiation is established by the complex interplay between transcriptional regulators, cis-regulatory elements and the chromatin landscape, all within the physical constraints imposed by 3D nuclear architecture^31–37^. This coordinated remodeling is not only limited to development, but is also important for neuronal reprogramming^38^ and disease^39,40^. However, it remains unclear if and how epigenetic mechanisms contribute to the progressive restriction of lineage potential in neural stem cells, to their ability to generate different type of projection neurons and to the switch to gliogenesis.

Here we characterized, in parallel, changes in the intrinsic transcriptional and epigenetic states of neural stem cells (NSC) and their differentiation fate biases across developmental time. We comprehensively profiled the transcriptional and epigenetic landscapes of mouse cortical development at single-cell resolution over a critical window in cortical development. Using differentiation flow models, we inferred NSC differentiation rates and specification preferences over time. We further dissected the underlying epigenetic mechanisms and defined the genome-wide interplay between chromosome accessibility, DNA methylation, 3D genome organization and enhancer activity in purified populations of NSCs across the same developmental window. Collectively, our data suggest that progressive epigenomic reprogramming of NSCs supports their dynamic differentiation fate biases over time and highlights novel mechanisms that promote the plasticity and flexibility of the NSC multipotent state. Inferred gene expression across metacell states can be viewed interactively at https://apps.tanaylab.com/MCV/mmcortex/.

## Results

### A Metacell flow model identifies lineage trajectories during mouse corticogenesis

To quantitatively and comprehensively understand self-renewal and diversification of NSCs during corticogenesis, we profiled the single-cell transcriptome and the chromatin accessibility landscape in parallel, sampling daily across a critical window of mouse cortical development (embryonic day E13 – E18, **Fig 1A**). Focusing first on the scRNA-seq data, which comprised of 42,172 cells across all timepoints, we inferred metacells and a manifold structure over them (**Fig 1B, S1A-B**)^41,42^. We corrected ambient noise using MCnoise (Methods), and excluded cell types of non-cortical origin, such as interneurons and microglia (Methods, **Fig S1C**).

**Figure 1:**
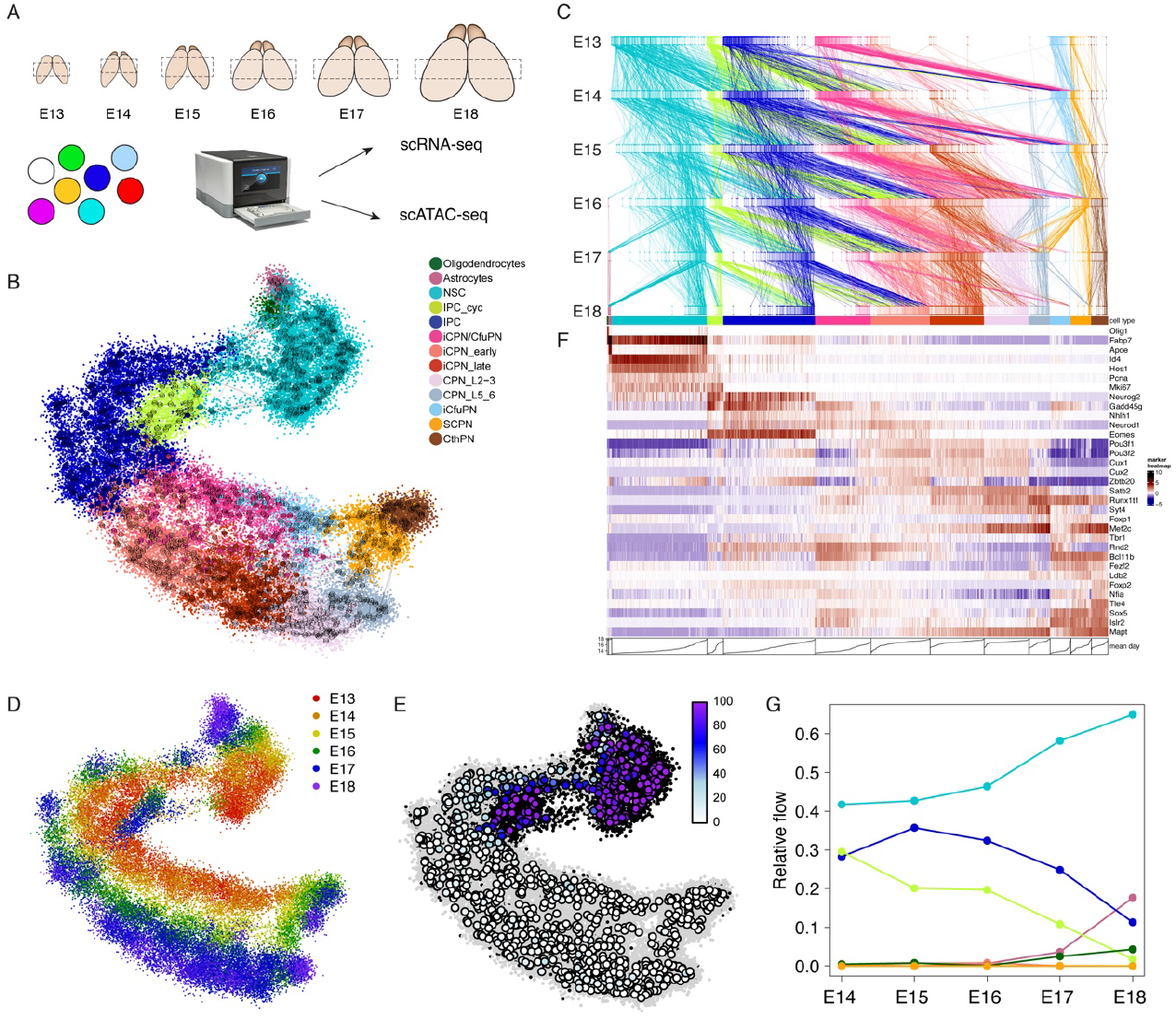
Comprehensive identification of lineage trajectories in mouse cortical development. A) Schematic representation of the single-cell experimental approach. Mouse somatosensory cortex was sampled at 24hr intervals between E13 and E18 and used for scRNA-seq and scATAC-seq assays. B) Two-dimensional UMAP projection of metacells (circles colored by cell type), edges of metacell similarity graph and single cells (dots colored by cell type). C) Visualization of metacell temporal flow model; vertical bars are metacell fractions in time points, diagonal lines represent flows between metacells. D) Single cells positioned by 2D projection of metacells and colored by time point. E) Metacells colored by cell cycle score, representing the percentage of cells classified as proliferating in each metacell. F) Heatmap of relative expression of marker genes across the metacell manifold. G) Relative temporal outflow from NSCs per time point (e.g. E13 to E14, E14 to E15, etc.), aggregated and colored by cell type.

Next, we inferred differentiation flows using the metacell flow algorithm ^43,44^ (**Fig 1C**, Methods). Flows link the observed cell ensemble distributions over time (**Fig 1D**) such that cells progress between metacells with maximal transcriptional similarity, with each cell producing progeny according to its metacell’s estimated proliferation rate (**Fig 1E, S1D**). In our data, proliferation rates were estimated to vary between 0 for postmitotic neurons and 2 doublings per 24h for NSCs^45^ (Methods).

We initially annotated metacells using known marker genes for NSCs, intermediate progenitor cells (IPCs), corticofugal (CfuPN) and callosal projection neuron (CPN) subtypes, astrocytes and oligodendrocyte precursors (OPCs) (**Fig 1F**). The inference of differentiation flows is agnostic to cell type annotation, allowing us to use flows for defining refined subsets of cycling IPCs, immature neurons (iCPN/CfuPN), corticofugal-biased immature neurons (iCfuPN) and callosal-biased early (iCPN_early) and late (iCPN_late) immature neurons (Methods). Overall, the output of the manifold and flow model are 1,093 annotated metacells, and inferred probability mass flows between them across the six time points.

When examining cell type fractions over time, we noticed that NSC, IPC/IPC_cyc and CfuPN (containing subcerebral – SCPN and corticothalamic – CthPN projection neurons) cell type fractions decreased (**Fig S1E**), while CPN proportion increased, consistent with the switch from deep layer to upper layer neurogenesis. Additionally, astrocytes and OPCs emerged in significant numbers at E17, indicative of the NSC fate shift from producing neurons to astrogliogenesis. To further capture the trends in proliferation and differentiation, we extracted the total probability mass flows aggregated over cell types at each time point (**Fig S1F**). NSC maintenance rates varied from 42% (E13->E14) to 62% (E17->E18), consistent with asymmetric-neurogenic proliferation of NSCs, whereby in each division one daughter cell remains an NSC, and another differentiates into an IPC. NSCs switched over time from a complete dominance of neurogenic IPC/IPC_cyc differentiation, to later onset of astrocyte and oligodendrocyte differentiation at E17 (**Fig 1G**)^2^.

### NSC fate bias changes with time and is synchronized with the cell cycle

To uncover the temporal dynamics of fate bias in stem cells, we first analyzed groups of genes co-varying between NSC metacells (**Fig S2A-B**). This identified gene clusters associated with the cell cycle, as well as genes that exhibit increased or decreased expression over time (Supplementary Table 1). To understand NSC fate predispositions and stemness signature, we also identified a neurogenesis/IPC-biased gene module and a gliogenesis/astrocyte-biased gene module (**Figs. 2A – top/middle**, Methods), to which we added genes enriched in NSCs relative to both astrocytes and IPCs, but not correlated with cell cycle phase, defining a core ‘stemness’ gene module (**Fig 2A – bottom**, Methods).

**Figure 2:**
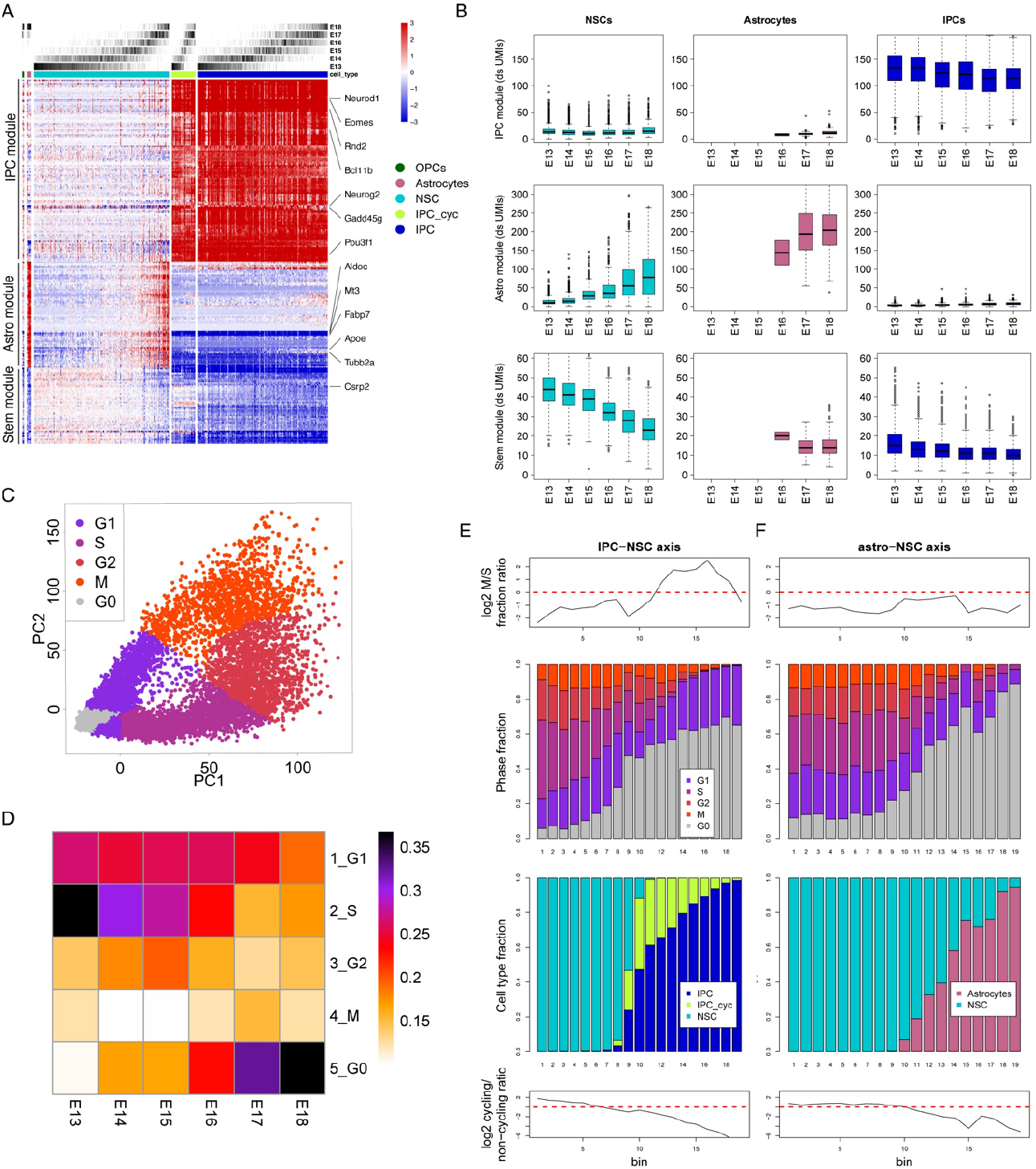
Temporal dynamics of NSC fate bias in the developing cortex. A) Gene expression across metacells, relative to mean expression in NSCs, of IPC, astrocyte and stem gene modules. Key marker genes are also labelled. B) Gene expression (total UMIs) of gene modules per cell type in downsampled single cells (3k UMIs/cell) per time point. C) First two principal components of the cell cycle in all cells in manifold, colored by inferred cell cycle stage. D) Fraction of NSCs in different cell cycle stages per time point. E) Ratio and scaled histogram of cell cycle phase and cell type fraction across NSC transition into IPC (left) or astrocytes (right). Single cells were divided into 20 bins in NSC->IPC and NSC->astrocyte trajectories. Top – log2 ratio of fraction of cells in M over S phase per bin; 2^nd^ from top – fractions in cell cycle stages per bin; 2^nd^ from bottom – cell types per bin; bottom – log2 ratio of cycling vs non-cycling cells per bin.

Next, we profiled the activity of IPC, astrocyte and stemness gene programs across cell types and over time (**Fig 2B**). We observed that the expression of the IPC program remained consistently low in NSCs across time, while the expression of astrocyte-linked gene increased by a factor of ∼8 between E13 and E18, reaching levels similar to those in astrocytes. In contrast, the stemness signature of NSCs decreased almost two-fold over time but remained at higher levels than in other cell types.

To investigate the link between differentiation fate and the cell cycle, we phased single cell profiles into G0, G1, S, G2, and M fractions based on gene expression (**Fig 2C, Fig S2C-E**, Methods). For IPCs and neuronal types, over 96% of the cells were assigned to the G0 (60.7%) or G1 (35.6%) phases (**Fig S2C**). For NSC, we observed slowing down of proliferation with time, with an increasing fraction of NSCs linked with the G0 stage (**Fig 2D**, increasing from 9% to 32%). These results are consistent with the lengthening of the NSC cell cycle observed during cortical development^19,46^.

To further dissect the relationship between cell cycle phase and fate, we analyzed the trajectory of NSC to IPC differentiation over 20 bins (**Fig 2E**). We observed relative enrichment of M-phase cells compared to S-phase cells (from bin 13-17), together with an overall convergence to a non-proliferating state (**Fig 2D, S2F**). This observation suggests a replication-first model for IPC differentiation in which the IPC program is activated primarily in G1. In contrast, grouping cells along the NSC to astrocyte trajectory did not reveal a similar trend (**Fig 2F**), consistent with a gradual convergence of the NSC toward the astrocyte program (**Fig S2F-I**).

In summary, our model suggests NSCs progressively lose their proliferative capacity, while gradually activating an astrocyte program, where there is possible synchronization of fate selection and the mitotic cycle.

### Distinct temporal trends of neural chromatin remodeling from a slowly shifting NSC state

Changes in NSC differentiation fates over time may be driven either by changing external signals, or by shifting cell intrinsic NSC state, or a combination of the two. To gain insight into the epigenomic signatures underlying the cell-intrinsic component of these shifting predispositions, we generated scATAC-seq libraries from 28,229 cells isolated from the same samples used for scRNA-seq (**Fig 1A, Fig S3A-C**, Methods). We developed an algorithm to map scATAC profiles over the scRNA-seq metacells using matching gene variation (Methods, **Fig S3D-E**). This resulted in an inferred accessibility score over metacells for ∼120k genomic hotspots (peaks). Among these, ∼19k peaks were proximal (<1kbp) to an annotated transcription start site (TSS), while the remaining ∼102k peaks were defined as TSS-distal, hereafter referred to as *CREs*.

Next, we grouped TSSs and CREs into 10 and 60 clusters, respectively, based on accessibility patterns across metacells (**Methods**). As expected, TSS clusters showed little variability across metacells (**Fig 3A**), consistent with previous results^31^. In contrast, CRE clusters were broadly separated into a constitutively accessible group (∼34k loci, **Fig 3B**, class I) and clusters with variable accessibility (classes II-VI). Within the variable groups, we identified two key groups associated with stem and progenitor cell activation. Specifically, class II CREs were accessible in NSC but lost activity immediately, or with some delay, upon transition to IPC, while class III CREs transiently gained accessibility in IPC, followed by either rapid or slow decline in iCPNs. Additional CRE clusters corresponded to regulatory elements active in more differentiated neuronal states: class IV was biased toward the corticofugal neural states (iCfuPN/SCPN/CthPN), class V toward CPN states and class VI displayed a late pan-neuronal activation pattern. Interestingly, analysis of motif enrichment in these CRE clusters (**Fig 3B-right**) uncovered a rich combinatorial structure, with intermediate progenitor and immature neurons characterized by a unique combination of motifs enriched in stem cells (NFIB, EMX1) and motifs specific to differentiated states (NeuroD/G, MEF2C).

**Figure 3:**
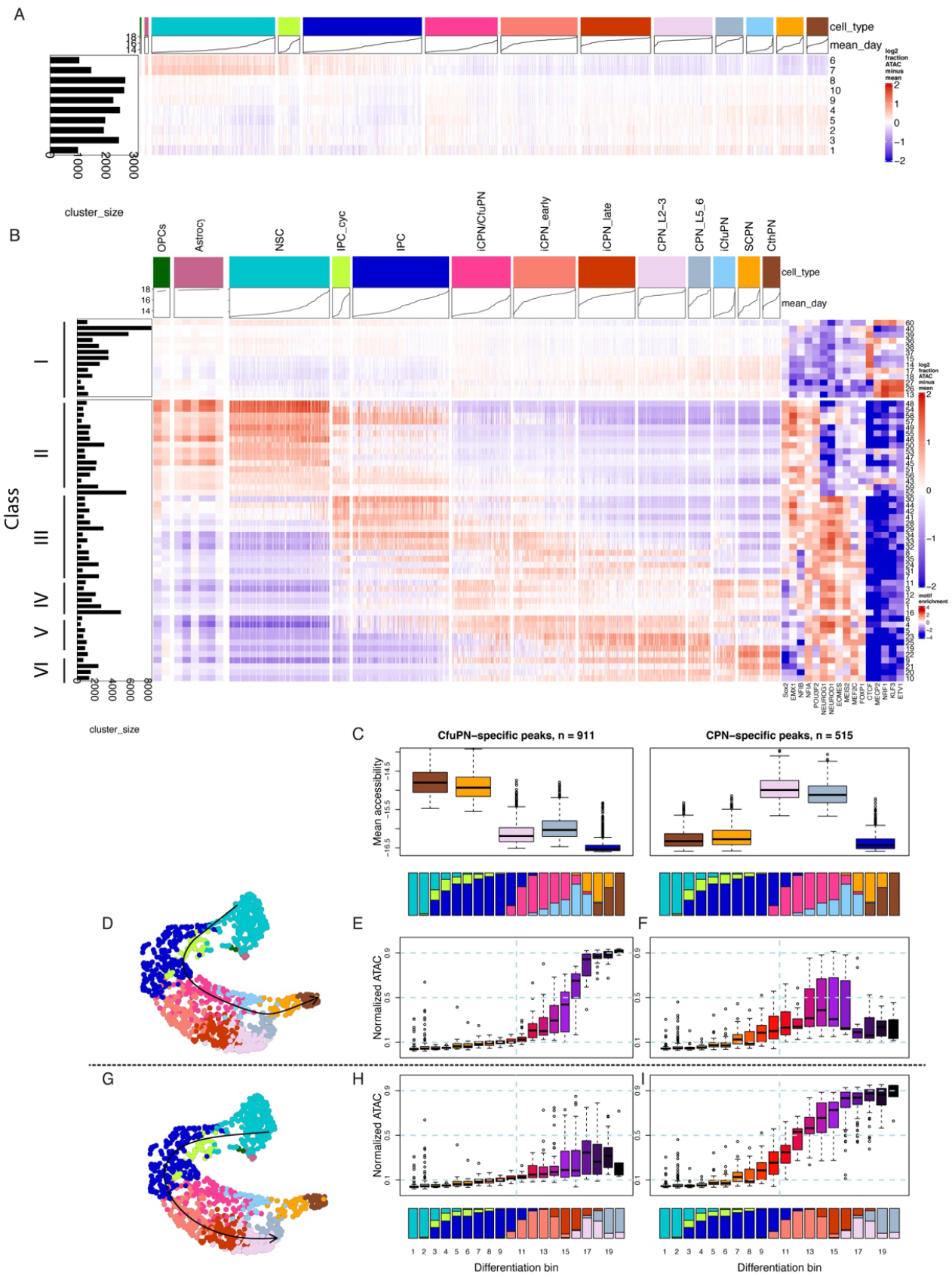
Widespread remodeling of the epigenetic landscape in projection neurons subtype trajectories. A) Heatmap of deviation from mean (log2 fraction) accessibility across metacells in TSS-proximal CRE clusters. Metacells are ordered by cell type (top, colors) and mean metacell day (top, line plots). Numbers on the left represent CRE cluster size. B) Same as A but for TSS-distal CRE clusters. Left – CRE cluster class. Right – heatmap of motif enrichment (log2 of observed over expected high-affinity sequences) for selected TF PWMs. C) Mean accessibility of CfuPN-(left) and CPN-specific (right) CREs across select cell types. D) Approximate CfuPN trajectory across the manifold. E) Mean normalized accessibility of CfuPN-specfic CREs across differentiation bins in the CfuPN trajectory. F) Mean normalized accessibility of CPN-specfic CREs across differentiation bins in the CfuPN trajectory. G) Approximate CPN trajectory across the manifold. H) Mean normalized accessibility of CfuPN-specfic CREs across differentiation bins in the CPN trajectory. I) Mean normalized accessibility of CPN-specfic CREs across differentiation bins in the CPN trajectory.

To dissect the accessibility landscape underlying the major cortical fate decision differentiating CPNs from CfuPNs lineages, we decoupled subtype specification from the overall pan-neuronal maturation process. We identified 911 CREs with CfuPN-specific late (i.e. low in IPC) activation and 515 CREs with specific activation in CPN (**Fig 3C – top boxplots**). We then traced the accessibility of these loci in their respective differentiation trajectories over the transcriptional and ATAC manifold (CfuPN trajectory – **Figs. 3D,E,F**, CPN trajectory – **Figs. 3G,H,I**). We found that CfuPN-specific CREs were activated late in the CfuPN trajectory (**Fig 3E**), with minimal representation in the immature states. In contrast, CPN-specific CREs were activated earlier in both trajectories (**Figs. 3F,I**), and subsequently suppressed in the later stages of the CfuPN trajectory (**Fig 3F**).

Overall, these results are consistent with postmitotic refinement of the two lineages, which is associated with epigenome rewiring predominantly in distal regulatory elements. Furthermore, our data indicate that repression of a CPN-associated epigenetic landscape in immature neurons (potentially mediated by the key repressive TFs of the CfuPN lineage such as Fezf2^47,48^ and Sox5^49^) plays an important role in the specification of projection neurons subtypes in the cortex.

### Accessibility gain in NSCs is accompanied by progressive loss of CpG methylation

We next extended our epigenomic profiling of NSC fate bias and performed bulk methyl-HiC^31^ on sorted Pax6^+^/Tbr2^-^ cells, corresponding to NSCs, from E13-E17 mouse embryonic somatosensory cortex (**Fig 4A**, Methods). This comprehensive profiling allowed us to quantify differential methylation at 99,078 well-covered TSS and CREs. As expected, accessible TSSs were almost invariably unmethylated (88% at m ≤ 0.05 (**Fig S4A**)). In contrast, constitutively accessible CREs (e.g. class I) showed more non-zero methylation (46% at m<0.05) and overall low methylation levels (81% at m ≤ 0.3 across all time points), whereas CREs with lineage specific accessibility exhibited more variable characteristics of NSC methylation (**Fig 4B**).

**Figure 4:**
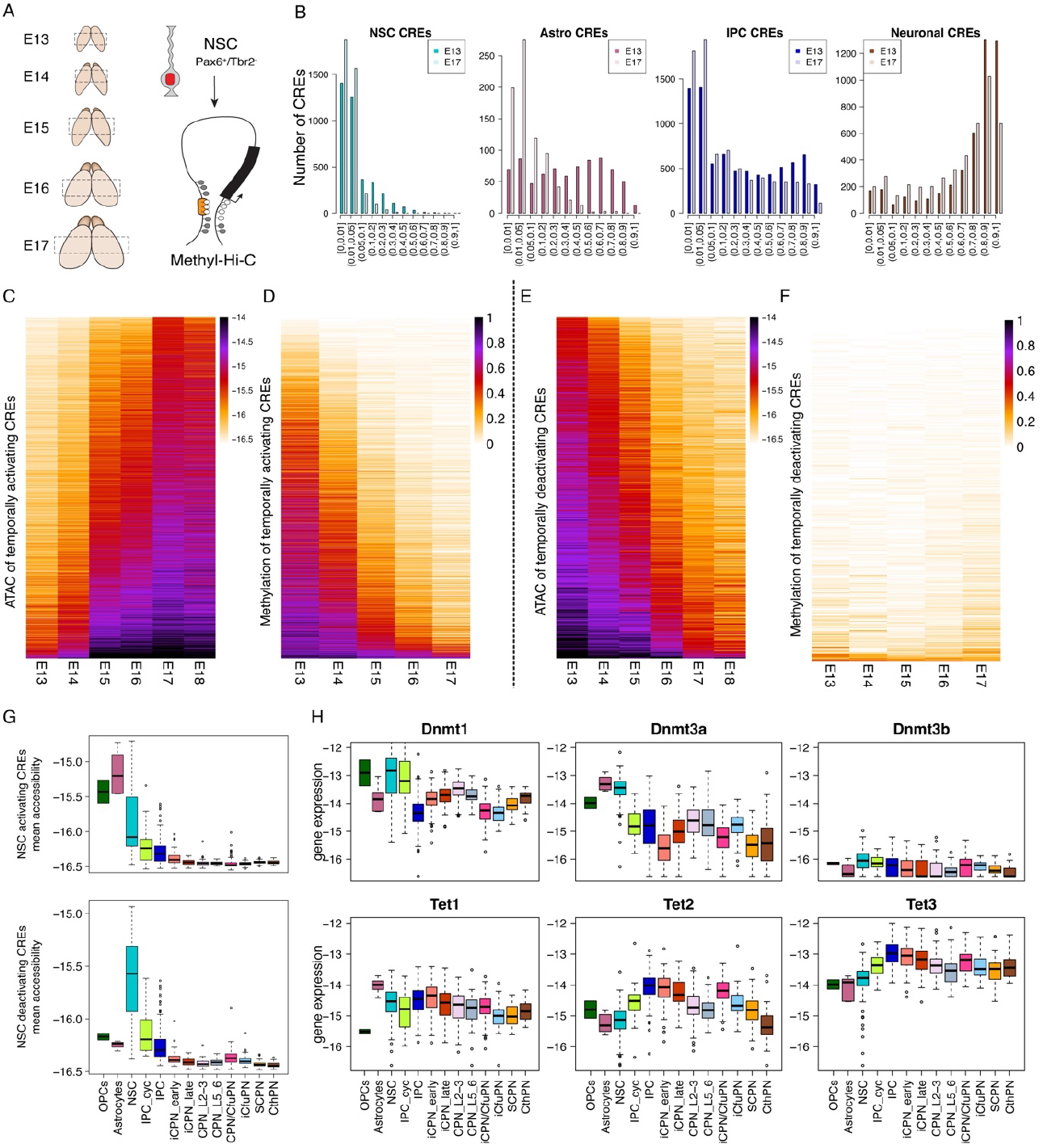
Distinct DNA methylation dynamics are associated with the changes in NSC temporal fate bias at CREs. A) Schematic representation of the experimental approach to profile 3D genome organization and DNA methylation in NSC. Somatosensory cortex was extracted from E13-E17 embryos. Dissociated NSCs were isolated by FACS and subjected to the methyl-Hi-C assay. B) Histogram, showing the number of cell type-specific CREs (as determined by the ATAC manifold) in methylation bins in the first (E13) and last (E17) time points assayed. C) Heatmap of chromatin accessibility of temporally activating CREs in NSCs. D) Heatmap of DNA methylation of temporally activating CREs in NSCs. E) Heatmap of chromatin accessibility of temporally deactivating CREs in NSCs. F) Heatmap of DNA methylation of temporally deactivating CREs in NSCs. G) Boxplots showing cell type-averaged accessibility of temporally activating (top) or deactivating (bottom) CREs in NSCs. H) Boxplots showing mean expression of DNA de/methylation enzyme genes in metacells by cell type.

NSC-specific CREs were predominantly un-(64% m<0.05) or lowly methylated (94.0-99.7% m< 0.3 across time points). Conversely, IPC CRE clusters included more sites with high methylation (m<0.3 only 57-71%), and most neuron specific CREs were highly methylated (m<0.3 14% to 22%). In general, CREs were biased towards methylation loss between E13 and E17 in NSCs (**Fig S4B**), while methylation in TSSs remained largely unchanged. Interestingly, astrocyte-specific CREs (defined based on activation exclusively upon astrocyte differentiation), showed remarkable loss of methylation in NSCs over time (from 43% m ≤ 0.3 at E13 to 95% at E17 – **Fig 4B**). These results suggest that loss of DNA methylation in NSC may be associated with priming and can participate in the regulation (and in particular, slow down) activation of the astrocytic program.

Accessibility and methylation are traditionally considered anti-correlated, a trend that we also observed in our data (**Fig S4C**). Consistent with this general pattern, CREs gaining accessibility over time in NSC frequently became demethylated indicating concomitant regulation (**Fig 4C,D**). In contrast, loci that lost accessibility in NSCs over time remained lowly methylated even at E17, suggesting that accessibility and methylation can be temporally decoupled within the same lineage (**Fig 4E,F**). Moreover, further analysis of cell type-averaged accessibility of temporally regulated CREs in NSCs revealed that regions that gained accessibility (“activating”) retained high accessibility only in glial types, while CREs that lost accessibility (“deactivating”) were NSC-specific (**Fig 4G**), and genome-wide accessibility of late NSCs was more similar to astrocytes than to early NSCs (**Fig S4E**).

To gain mechanistic insights into this unique pattern of temporal regulation, we examined the expression of different enzymes associated with DNA methylation. We found that NSCs expressed the de-novo methyltransferase Dnmt3a at relatively high levels (**Fig 4H**), which are rapidly repressed upon differentiation to IPCs, but sustained in glial cell types. Furthermore, the demethylation factors Tet2,3 were expressed at low levels in NSC and became induced during neuronal differentiation, while Tet1 was robustly induced upon differentiation to astrocytes (**Fig 4H**). Among NSC metacells, the mean trend of accessibility at “activating” CREs was weakly (r = 0.16) yet significantly (p=0.02) correlated with Tet1 expression (but not Tet2/3), while accessibility at “deactivating” CREs weakly but significantly correlated with Dnmt1/3a/3b levels (r = 0.18, 0.13, 0.23; p = 0.008, 0.058, 0.001) (**Fig S4D**). These relatively modest changes could be further reinforced by the dynamic expression of co-factors and protein regulators that modulate DNA methylation patterns over time in NSC. Indeed, we identified Sall3, a transcription factor previously shown to interact with DNMT3A and inhibit CpG methylation^50^. Despite having no known role in the cerebral cortex, Sall3 was temporally regulated and became expressed in late NSC and astrocytes (**Fig S4F**).

Collectively, these data suggest that NSC fate bias is correlated with, and perhaps influenced by, dynamics of the DNA methylation machinery, including targeted demethylation at astrocytic regulatory elements that are progressively remodeled toward activation in late neurogenesis in NSCs.

### Hi-C uncovers 3D chromosomal remodeling in NSCs over time

To compare the epigenome dynamics at the linear genome level with changes in higher-order chromatin structure, we next examined 3D genome organization, focusing on chromatin insulation^37^ (Methods). Overall, insulation scores were highly conserved between time points (**Fig S5A**). Nevertheless, we identified 335 genomic regions with insulation changes between E13 and E17 (**Fig 5A**, Methods). Among these, 100 regions were characterized by increased insulation over time (“insulating”), while 235 lost chromatin insulation (“*de-insulating”*) (**Fig 5A**). Examples of de-insulating (Hapln2) and insulating (Tnc) loci are shown in **Fig 5B-G**: *Hapln1* is positioned at the border of two TADs (**Fig 5B,C**) and is not expressed robustly in NSCs (**Fig 5D**); the gene locus undergoes gradual compaction (**Fig 5C**), leading to decreased insulation score (**Fig 5B**), prior to its expression in astrocytes (**Fig 5D**). Similarly, the Tnc locus is positioned at the edge of a TAD that is partially in contact with a neighboring TAD (**Fig 5E,F**); the TAD containing *Tnc* is increasingly compacted (**Fig 5F**) along with increased *Tnc* expression, reaching peak expression in astrocytes (**Fig 5G**). Analysis of CRE distribution around insulators showed that loss of insulation was correlated with CRE activity (significant enrichment within 100kb around the site, p<5e-4, Methods), while increasing insulation was observed in regions with relative depletion of active CREs (**Fig 5H**).

**Figure 5:**
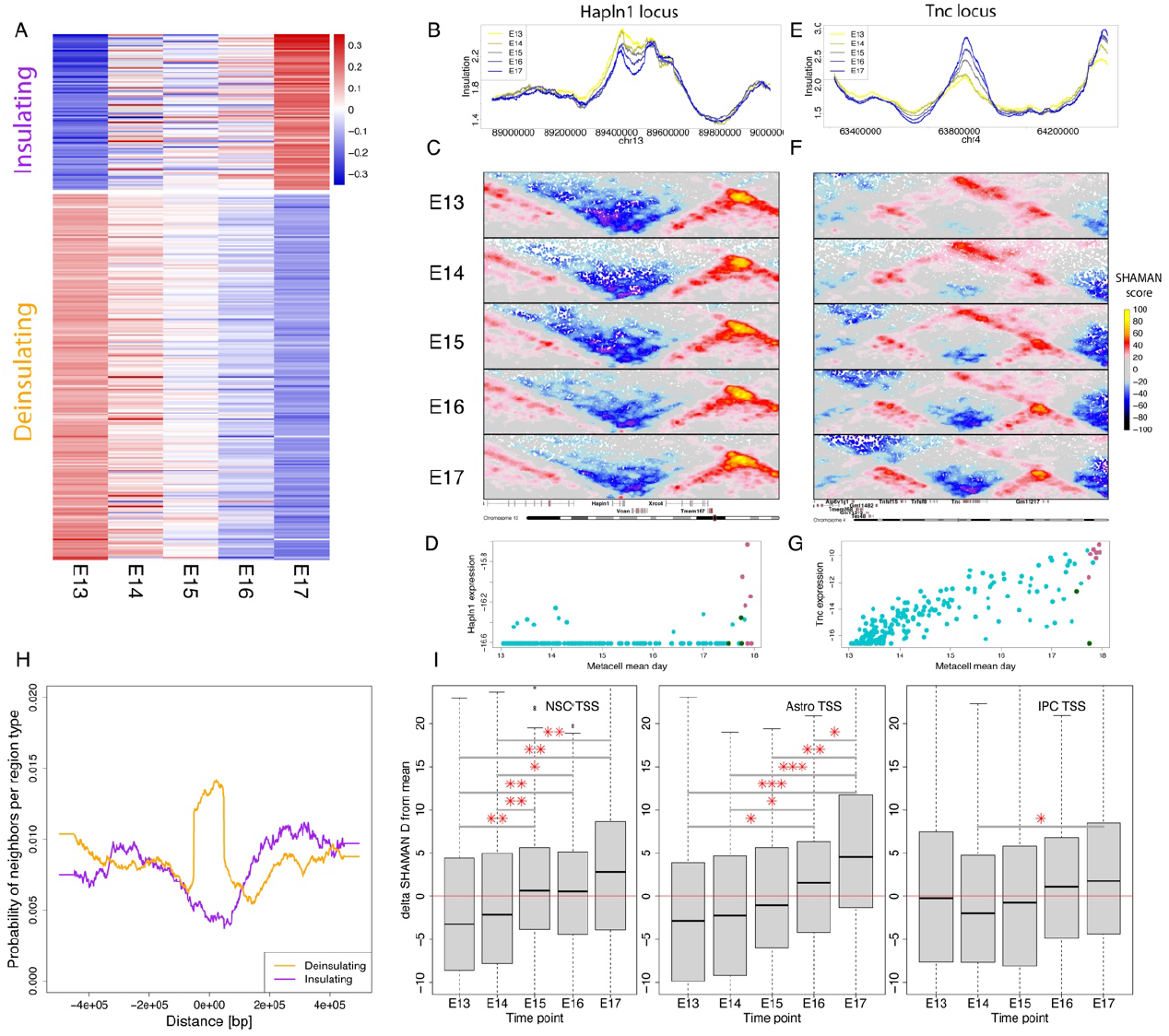
3D epigenome rewiring suggests potential CRE cooperativity in NSC epigenome remodeling. A) Heatmap showing deviation from mean insulation score in regions with increasing (insulating) or decreasing (deinsulating) insulation scores in NSCs across time. B) Insulation score across time in ∼1Mbp window around the *Hapln1* gene. C) Normalized interaction frequencies (SHAMAN scores) in same window as C) across time. D) Hapln1 expression in NSC, astrocyte and OPC metacells across time. E-G) Same as B-D but for the *Tnc* gene. E) Rolling average (window size = 60kbp) probability of neighboring accessible CREs in NSCs around insulating (purple) and deinsulating (orange) regions. F) Deviation from mean SHAMAN scores (D) between CREs that are temporally activating in NSCs and TSSs of cell type-specific genes.

We next focused on the chromosomal contacts of CREs that gain accessibility over time in NSCs. We observed that these CREs engaged in significantly stronger interactions with Astrocyte TSSs (as well as NSC TSSs) across multiple time points (**Fig 5I**), while exhibiting weak or non-significant contacts with IPC TSSs.

Collectively, our data on both insulation dynamics and CRE contacts suggest that, in addition to gradual changes in DNA methylation, NSCs chromosomal architecture evolves over time to form a potentially astrocytic-biased architecture. This is accompanied by decreased potential for generating IPCs and an increased probability for conversion toward astrocytes, potentially contributing to the fate switch of NSC at the end of neurogenesis.

### Modelling IPC chromatin accessibility using TF affinities and epigenomics

In order to characterize the interplay among cis-regulatory and epigenetic mechanisms guiding NSC differentiation, we developed a machine-learning model to capture the dynamics of NSC-to-IPC chromatin accessibility remodeling. First, we inferred specificities for 16 candidate motifs from CRE sequences (**Fig S6A**, Methods). These motif-specific features were then integrated with further parameters defining the epigenetic baseline state, including mean NSC accessibility and methylation level at each CRE. In addition, we incorporated regional activity metrics by summing up accessibility (within a 50 kbp window) and transcription (within a 0.5 Mbp window) around each CRE locus.

Based on these features, our model achieved surprising accuracy, with an R^2^ of 0.62 on held-out data (**Fig 6A**), and an R^2^ of 0.41 when applied to a generic, non-cortex-related set of CREs (**Fig S6B**, Methods). The most important model features (based on SHAP analysis, **Fig 6B**), were the baseline NSC ATAC level, an E-box motif (potentially linked to bHLH TFs such as NEUROG/NEUROD), a T-box motif (potentially associated with EOMES) and mean NSC methylation level. These top features, however, provided only around half of the total SHAP values, with additional features such as Fos::Jun and SOX motifs^31^, contributing to the prediction of the IPC-NSC accessibility difference.

**Figure 6:**
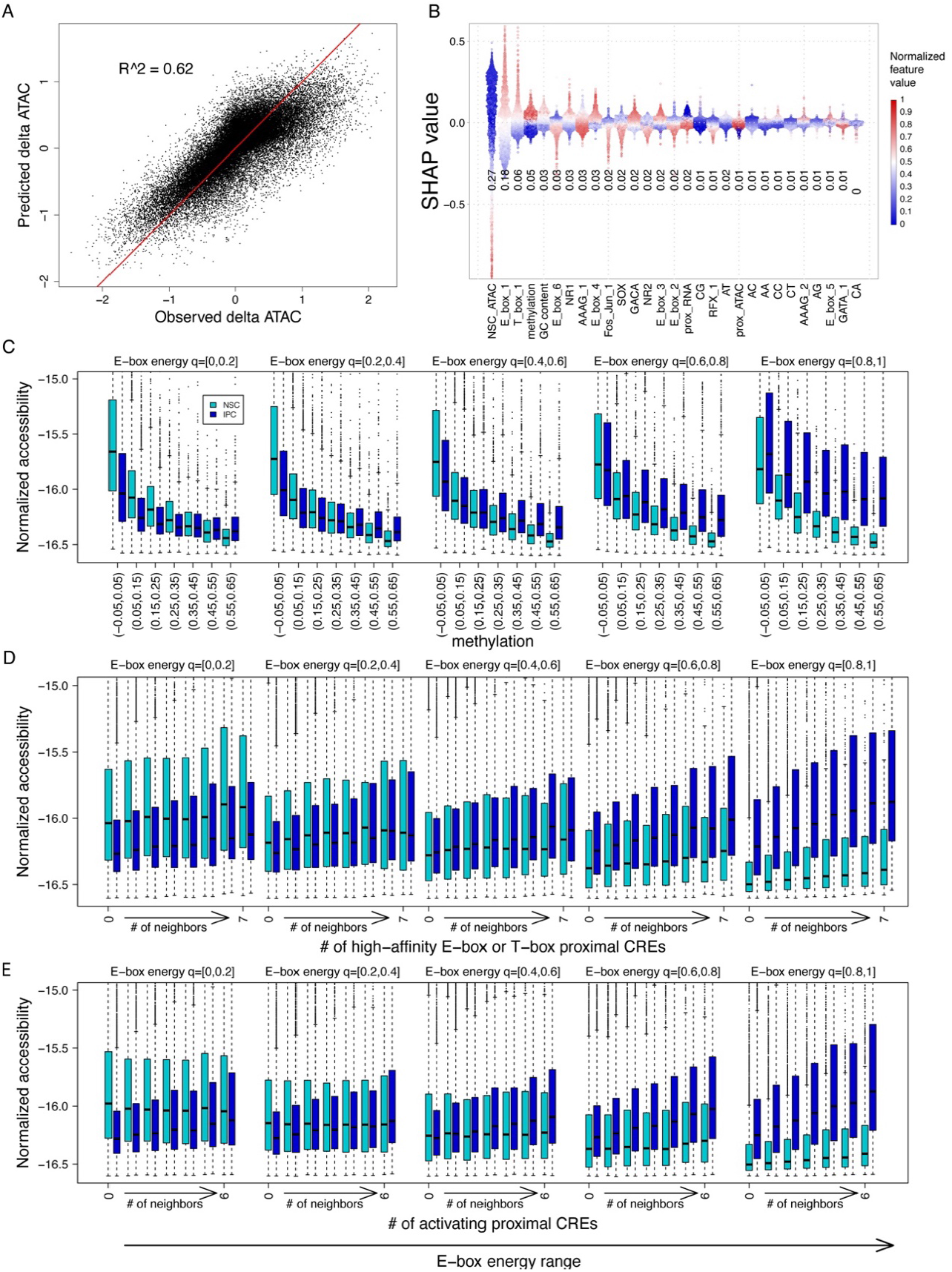
Machine learning model quantifies the contribution of TF affinities and epigenome in NSC lineage transitions. A) Scatterplot of mean IPC vs mean NSC chromatin accessibility as predicted by the xgboost sequence and epigenomic model vs. observed difference. B) Distribution of SHAP values for main features of xgboost model. Text labels are the fraction of the sum of each feature’s absolute value SHAP from the total absolute value SHAP across all features. Points are colored by the linearized feature value. C) Mean NSC and IPC accessibility for all TSS-distal CREs, stratified by E-box energy and methylation. D) Same as C but stratified by, instead of methylation, the number of neighboring CREs with high E-box/T-box affinity. E) Same as C but stratified by, instead of methylation, the number of neighboring CREs that are accessible in NSC and/or IPC.

Understanding the mechanism underlying a prediction necessitates careful stratification of the model’s features, which are in many cases highly co-dependent. This is specifically important when the model considers a set of CREs that are pre-selected to be active in either NSCs or their progenies. For example, the E-box motif was highly correlated with NSC methylation levels in this CRE set (**Fig S6D**), but this effect was due to the enrichment of E-box elements in IPC-specific CREs, that were methylated in NSC prior to their activation. Only when correcting for E-box affinity by stratification, a negative correlation between DNA methylation and IPC accessibility (**Fig 6C**) was observed, consistent with the model suggested earlier (**Fig 4)**.

Finally, we asked if the local epigenetic context affects the activity of lineage-specific CREs. Analysis of local CRE E-box sequence affinity conditioned on the presence of proximal CREs with high E-box and/or T-box sequence affinities within a 20kbp radius (**Fig 6D**) corroborated our previous findings (**Fig 5**). Specifically, CREs with medium-high, or high E-box affinity were further activated in the vicinity of additional high-affinity CREs. This effect was found to be most pronounced when considering neighboring elements within 20kbp (**Fig S6E**). Moreover, the number of active neighboring elements, without filtering for their sequence content, was also predictive of higher accessibility in the IPC state (**Fig 6E**).

Overall, these results point to a complex and multivariate regulation to lineage transitions in the cortex. While the initial epigenetic state and specific transcription factor motifs play predominant roles, the local epigenetic context and synergistic interactions among multiple CREs further contribute to ensure robust cell-fate transitions.

### *In vivo* reporter assays decouple autonomous from context-dependent CRE regulation

To validate and further dissect autonomous vs. context-dependent CRE regulation, we selected sequences of 11,905 loci and subjected them to an *in vivo* massively parallel reporter assay (MPRA) as previously published^31^ (Methods). Episomal libraries were electroporated into embryonic cortices between E12 and E16, and NSC and IPC cell populations were isolated via FACS 24hr after each injection (**Fig 7A**, Methods). The resulting sequencing libraries were processed using MPRAflow^51^ and scored using MPRAnalyze^52^, revealing that 9,101 CRSs (76%) were found to contribute significantly to reporter expression in at least one time point.

**Figure 7:**
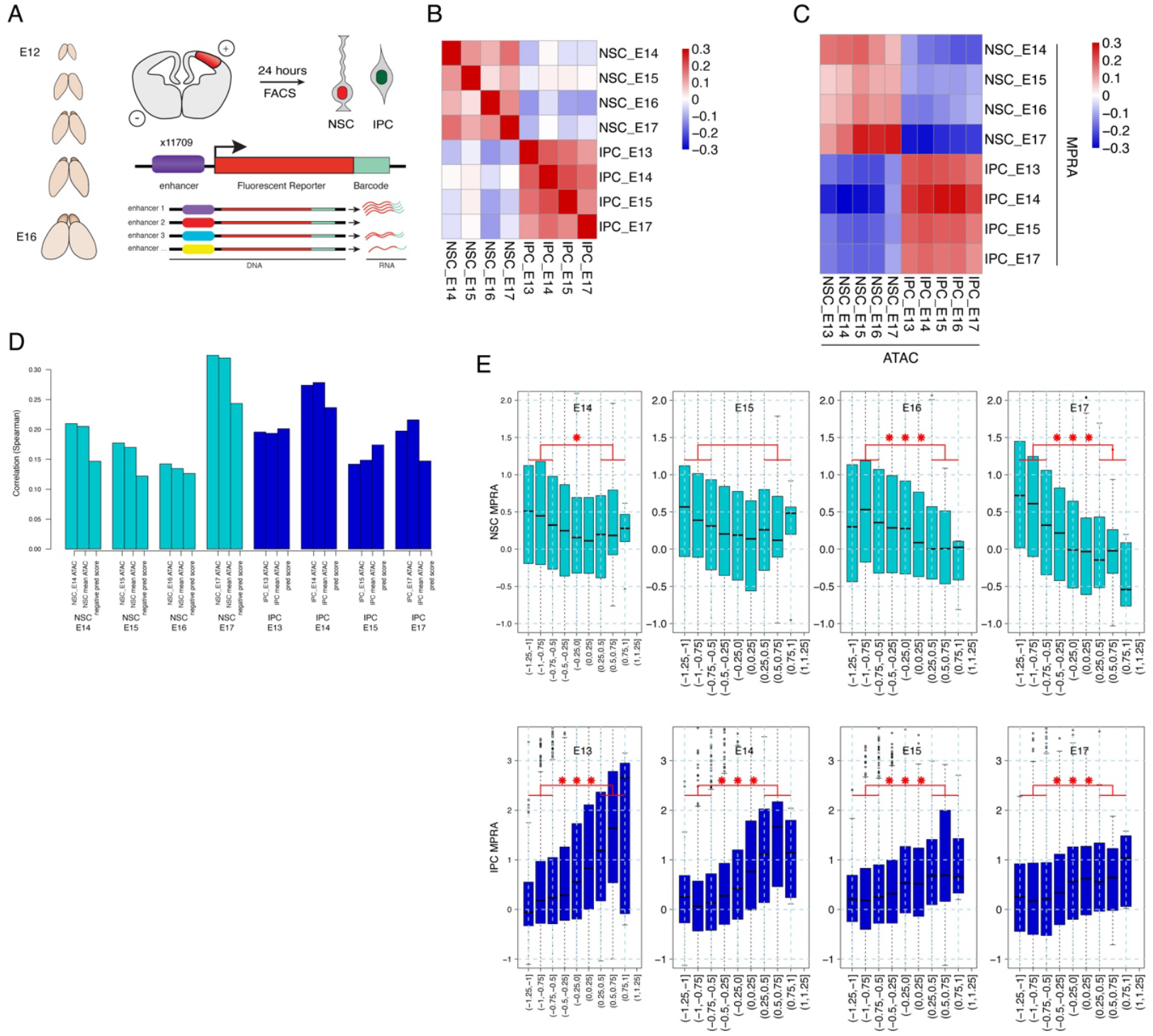
Massively parallel reporter assays *in vivo* decouple context-dependent from autonomous CRE activation. A) Schematic representation of the *in vivo* MPRA experiment. B) Heatmap of correlations between normalized MPRA scores across time points and cell types. C) Heatmap of correlations between normalized MPRA scores (rows) and chromatin accessibility (ATAC) of cognate CREs in respective cell types (columns). D) Barplots depicting the correlation between MPRA scores and chromatin accessibility from same time point and cell type, mean ATAC of the respective cell type across all time points, and predicted score from xgboost sequence model. E) MPRA scores vs binned predicted sequence score by cell type and time point. Bars and asterisks denote pooled bins that were compared and significance of two-sided KS p-value (* < 5e-2, ** < 5e-3, *** < 5e-3), respectively.

After normalization and exclusion of two libraries (Methods), we observed distinct activity patterns in NSC and IPC across different sampling days (**Fig 7B**). Cross-correlation of chromatin accessibility with reporter activity showed that sequences associated with differential accessibility in NSC or IPC within their native chromosomal context, retained their cell type-specific activator autonomously and out of such context (**Fig 7C**).

Next, we adapted our IPC-NSC accessibility prediction model to rely on sequence alone, by fixing all epigenetic parameters to match an unmethylated and open context as provided by the reporter system (Methods). We then predicted an IPC-NSC activity score for each experimentally active sequence. This predicted activity score correlated with the MPRA signal at levels comparable to the those using time point-specific and cell type-averaged accessibility (**Fig 7D**). Moreover, stratification of profiled elements by their predicted NSC vs. IPC sequence potential (as in **Fig 6**) validated the model by showing scores are positively correlated with IPC activity, and negatively correlated with NSC activity, at all time points (**Fig 7E**).

Taken together, *in vivo* reporter assays validate our CRE sequence model and further highlight the distinction between inherent, autonomous sequence potential of CREs and their potential for synergistic activation within a chromosomal context. Notably, the temporal shift in NSC CRE regulation indicates that the fate bias change towards astrocytic identity is driven by changes in trans-factors’ activity, occurring in parallel with gradual NSC epigenetic programming (i.e. DNA methylation and chromosomal conformation).

## Discussion

In this manuscript, we dissected the process of neuronal stem cell differentiation and fate determination from embryonic day E13 to E18, an essential window of embryonic corticogenesis in the mouse. Our analysis combined single-cell gene expression and chromatin accessibility data, together with cell type-specific, genome-wide profiling of 3D genome organization, DNA methylation and activity of 11905 CREs, representing one of the most comprehensive datasets characterizing mammalian cortical development. Standardized sampling over time enabled us to infer the temporal differentiation dynamics from NSCs towards more differentiated progenitor states, distinct projection neuron subtypes and glia. Single-cell analysis, facilitated by metacell flow modeling, revealed that NSC gradually lose their transcriptomic stem cell signature, while converging toward an astrocytic program by E17. In addition, we demonstrated that the NSC mitotic cycle is synchronized specifically with IPC regulatory activation, an effect not observed during the NSC-to-astrocyte transition.

Our temporal model also allowed us to investigate intrinsic epigenetic factors underlying NSC fate bias. Although extrinsic factors, such as inter-cellular signals, niche structure and morphology^22^, undoubtedly contribute to fate decisions, our analysis illuminates key aspects of inherent NSC potential and the intrinsic mechanisms driving its progressive lineage restriction over time.

Across multiple scales and modalities, we demonstrated that corticogenesis involves extensive cell type-specific remodeling of CREs, in concert with the increased activity of specific TFs that regulate stem cell differentiation. Beyond the canonical differentiation trajectories between cell types, our temporally resolved analysis revealed gradual epigenetic remodeling, over absolute time, *within* cell types. In NSCs, temporal changes in chromatin accessibility, reflected by progressive gains or losses in CRE activity, were inversely correlated with DNA methylation dynamics. However, loss of accessibility in early NSC CREs was not accompanied by a corresponding increase in DNA methylation. Whether alternative repressive mechanisms (such as Polycomb-mediated repression^19,20^) are associated with such loci remains unclear. It is tempting to speculate that CRE repression during corticogenesis is governed by mechanisms and epigenetic dynamics distinct from those associated with CRE activation.

In addition to changes in the linear epigenetic landscape, we also observed significant dynamics in 3D genome organization. Chromosomal contact analysis revealed that NSCs gradually remodel specific TAD insulation anchors, a process that correlates with local CRE activity and transcriptional output. Furthermore, interactions between NSC CREs and astrocyte-specific TSSs become stronger prior to the switch to astrogliogenesis, suggesting that epigenetic priming within the regulatory landscape may facilitate this fate transition. Collectively, we have shown, across multiple modes of genomic regulation, that NSCs epigenome gradually drifts toward astrocyte state over time, offering insights into the mechanistic linkage between intrinsic epigenetic remodeling and evolving NSC fate bias.

Whether the epigenetic remodeling we observed represents a defining characteristic of NSC’s restricted differentiation potential remains an open question. Consistent with this notion, both Polycomb group and high-mobility group proteins have been shown to play an instructive role in NSC lineage progression and the switch to astrogliogenesis^53,54^. Our mechanistic modeling of CRE activity, enabled by the comprehensive and multimodal nature of our data and a novel machine-learning approach, provides critical insights into this process. We identified key TF motifs as strong predictors of chromatin accessibility dynamics during the NSC-to-IPC transition, yet other epigenetic features such as DNA methylation and chromosomal context (total activation potential of the regions surrounding the CRE) contribute further to this process. Notably, these features align with the epigenetic dynamics we directly profiled using methyl-Hi-C, highlighting their previously unrecognized roles in the temporal regulation of cortical development.

Finally, we further validated and assessed the autonomous capacity of CRE sequences to respond to TFs by performing cell-type specific MPRA on more than 11000 sequences across five developmental time days *in vivo*. This dataset confirms the influence of extrinsic factors relaying signaling cues through TFs and into the genome. Moreover, it highlights the distinction between inherent, autonomous sequence regulatory potential of CREs and their synergistic activation within a chromosomal context – reconciling a long-standing debate regarding the relative importance of intrinsic versus extrinsic cue in the context of NSC lineage specification.

Transcriptional atlases provide a crucial blueprint for understanding tissue developmental dynamics. Our data show that inference of such dynamics can rely on careful single-cell temporal sampling, metacell-based differentiation flow modelling and in-depth single-cell and population-averaged epigenetic profiling. In the future, modeling of tissue development would be further enhanced by spatio-temporal profiling, that can augment the observed temporal behaviors with an essential context describing inter-cellular signals and interactions. Going beyond single-cell trajectories (e.g. pseudo time over transcriptional manifolds), and into models that describe cell dynamics and are aware of epigenetic history and tissue context, will be essential for understanding the transition between stem cell multipotency and self-renewal and the formation of organized, balanced and functional brain, or any other tissue.

## Author Contributions and Notes

N. and S.V. performed experiments. Y.S., F.C., A.T. and B.B. analysed the data. A.L. established the MCView interface. B.B. and A.T. supervised the project. Y.S., A.T. and B.B. wrote the manuscript with input from all authors.

### Acknowledgments

Sequencing was performed at the Helmholtz Munich by the NGS-Core Facility and cell sorting was done at the Flow Cytometry Core Facility at the Biomedical Center, Ludwig-Maximilians-Universität. Work in the group of B.B. was supported by the ERA-NET Neuron (MOSAIC) and European Research Council Consolidator Grant (EpiCortex, 101044469). Work in the group of A.T. was supported by a European Research Council Advanced Grant (Cells2Tissues), and by the Israeli Science Foundation. Y.S. would like to thank all Tanay lab members for technical help and useful discussions.

## Materials and Methods

### Experimental model

Time-mated pregnant C57BL/6JRj mice were purchased from Janvier Laboratories and housed under standard conditions in compliance with local regulations set by the Regierung Oberbayern, Germany. Mouse embryos were used irrespective of sex. All experiments were conducted in accordance with national guidelines and approved by local authorities (Regierung Oberbayern, Germany; ROB-55.2-2532.Vet_02-19-175).

### scRNA-seq and scATAC-seq

Somatosensory cortices were isolated from E13, E14, E15, E16, and E17 embryos after removal of the meninges and dissociated using a papain-based neural dissociation kit (Miltenyi Biotec, Cat. N.: 130-092-628) following the manufacturer’s protocol. Library preparation for scRNA-seq (v3, 10x Genomics) and scATAC-seq (v1.1, 10x Genomics) was performed according to the manufacturer’s instructions, targeting a recovery of 6,000 nuclei per sample.

### Fluorescence-activated cell sorting (FACS)

FACS was performed as previously described^31^, following intracellular staining. A detailed protocol for the immunoFACS is available at https://www.protocols.io/view/immunofacs-b2a2qage/. In brief, dissociated cells were fixed for 10 minutes at room temperature using 1% freshly prepared Formaldehyde in PBS (ThermoFisher, Cat. N: 28908) under slow rotation, and the reaction was quenched by adding Glycine (Invitrogen, Cat. N: 15527013) to reach a final concentration of 0.2 M. Following fixation, the cells were centrifuged at 500g for 5 minutes at 4°C, washed once with a buffer composed of 1% BSA and 0.1% RNAsin plus RNase inhibitor (Promega, Cat. N: N261A) in PBS, and then incubated for 10 minutes at 4°C in a permeabilization solution containing 0.1% freshly prepared Saponin (Sigma-Aldrich, Cat. N: SAE0073), 0.2% BSA (ThermoFisher, Cat. N: 15260-037), and 0.1% RNAsin plus RNase inhibitor in PBS. After permeabilization, the buffer was removed by centrifuging the cells at 2500g for 5 minutes at 4°C, followed by staining with antibodies against Pax6 (1:40; BD Bioscience, Cat. N: 561664), Eomes (1:33; BD Bioscience, Cat. N: 566749), and Tubb3 (1:14; BD Bioscience, Cat. N: 560394) in a staining buffer (0.1% saponin, 1% BSA, 0.1% RNAsin plus RNase inhibitor in PBS) for 1 hour at 4°C under slow rotation. The cells were subsequently washed twice with the permeabilization buffer, once with a wash buffer containing DAPI (1:1000; ThermoFisher, Cat. N: 62248), and finally with wash buffer without DAPI. Each washing step involved a 5-minute incubation at 4°C under slow rotation, and cells were centrifuged at 2500g for 5 minutes at 4°C between washes. After the final wash, cells were resuspended in PBS with 1% BSA and 1% RNAsin plus RNase inhibitor, filtered through a 40µM cell strainer (ThermoFisher, Cat. N: 15342931), and promptly subjected to FAC-sorting. Sorted cells were either directly used for nucleotide isolation (MPRA) or flash frozen and stored at -80°C (Methyl-Hi-C and *in situ* Hi-C).

### Methyl-Hi-C

Somatosensory cortices were isolated from E13, E14, E15, E16, and E17 embryos after removal of the meninges and dissociated using a papain-based neural dissociation kit (Miltenyi Biotec, Cat. N.: 130-092-628) following the manufacturer’s protocol. Each biological replicate represents a pool of 4-6 littermates from separate mothers. Methyl-Hi-C and low-input *in situ* Hi-C on the sorted NSC were conducted as previously reported^31^ and the detailed experimental procedure can be found at https://www.protocols.io/view/methylhic-bif2kbqe/ and https://www.protocols.io/view/in-situ-hi-c-brd4m28w/, respectively. Briefly, frozen pellets of fixed cells were thawed on ice and then lysed using 0.2% Igepal-CA630 (Sigma-Aldrich, Cat. N: I3021). The cells were subsequently permeabilized with 0.5% SDS (Invitrogen, Cat. N: AM9823) and digested overnight at 37°C using 200U DpnII (New England Biolabs, Cat. N: R0543). Next, the sticky ends were filled in by incubating the nuclei for 4 hours at room temperature with DNA Polymerase I (New England Biolabs, Cat. N: M0210) in the presence of a nucleotide mix containing biotin-14-dATP (Life Technologies, Cat. N: 195245016) in DpnII buffer, followed by a proximity ligation step of at least 6 hours at 16°C. Afterward, the nuclei were lysed, and the proximity ligated DNA underwent reverse-crosslinking overnight at 68°C, was purified by ethanol precipitation, and then sheared to ∼550 bp fragments using a Covaris S220 sonicator. An end-repair was performed after sonication by incubating the samples with T4 DNA Polymerase (New England Biolabs, Cat. N: M0203) for 4 hours at 20°C. Before bisulfite conversion, approximately 0.01% of sheared and biotinylated fully methylated pUC19 (Zymo Research, Cat. N: D5017) and unmethylated lambda DNA (Promega, Cat. N: D1521) was added to the samples. Bisulfite conversion was then carried out using the EZ DNA Methylation-Gold kit (Zymo Research, Cat. N: D5005) followed by library construction with the Accel-NGS® Methyl-Seq DNA Library kit (Swift Bioscience, Cat. N: 30024) according to the manufacturer’s instructions up to the adapter ligation step. Afterward, biotin pulldown was performed using MyOne Streptavidin T1 beads (ThermoFisher, Cat. N: 65602), with five washes using a buffer containing 0.05% Tween-20 (Sigma-Aldrich, Cat. N: P9416) and two additional washes with low-TE water. Finally, libraries were amplified on the streptavidin beads using EpiMark Hot Start Taq (New England Biolabs, Cat. N: M0490) along with Methyl-Seq Indexing Primers (Swift Bioscience, Cat. N: 36024) using the following program: 95°C for 30 seconds; then 14 cycles of 95°C for 15 seconds, 61°C for 30 seconds, and 68°C for 60 seconds; followed by 68°C for 5 minutes; and a final hold at 10°C. The streptavidin T1 beads were then magnetically pelleted, and the libraries present in the supernatant were purified with 0.6x AMPure XP beads (Agencourt, Cat. N: A63881) to achieve an average fragment size of approximately 500 bp.

### Massive parallel reporter assay (MPRA)

The MPRA design followed the approach previously described by^31^, with the following specifications. The MPRA plasmid pool included 997 scrambled control sequences, which had matched GC content and were pre-screened to minimize the presence of expressed TF motifs, and 11,905 putative cis-regulatory sequences (CRSs) derived from CREs with temporally increasing, decreasing or stable accessibility patterns in NSCs, IPCs or neurons (roughly 1k per cell type and pattern combination). All sequences were centred on the accessibility peak and resized to 266bp.

A detailed protocol for MPRA plasmid pool generation is available at https://www.protocols.io/view/mpra-plasmid-pool-preparation-bxchpit6/. We electroporated the MPRA library into the cortices of mouse embryos at E12, E13, E14, E15 and E16 and collected the embryos 24 hours after. We then removed the meninges, dissected the somatosensory cortex and dissociated the cells using a papain-based neural dissociation kit (Miltenyi Biotec, Cat. N.: 130-092-628) according to the manufacturer’s protocol. Dissociated cells were stained for Pax6, Tbr2 and Tubb3, and specific populations were sorted using FACS as described above.

### *In-utero* electroporation

In-utero electroporation was carried out according to the previously reported protocol^31^. In summary, pregnant mice with time-mated plugs were anesthetized with isoflurane, the uterus was exposed, and 1–3 µl of plasmid DNA in PBS was injected into the telencephalic lumen. This injection was immediately followed by five pulses at 35 V, each lasting 100 ms with 1-second intervals, delivered via platinum electrodes (NepaGene, Cat. N: CUY650P1). At either 24 or 48 hours post-electroporation, the pregnant females were sacrificed, the embryos’ brains were dissected, and the electroporation efficiency was evaluated using a fluorescence stereomicroscope.

### scRNA-seq mapping and filtering

scRNA-seq data was processed using a standard CellRanger pipeline (version 3.1.0) to generate count matrices per library. Raw count matrices were then filtered by removing cells if they had more than 15% of UMIs mapping to the mitochondrial genome, or 0.5% of reads mapping to hemoglobin genes, or less than 1500 UMIs or more than 30,000 UMIs.

Doublets were removed using Doubletfinder^55^ with 30 principal components, log-normalization of gene expression (scale 10,000), 2,000 genes and clustering resolution of 0.6. A linear model was fit to 10x Genomics data on expected doublet fractions in proportion to cells loaded in a chip, and 0.75 times the expected fraction per sample was used a parameter for the calculation of the Poisson exponent in the Doubletfinder pipeline. The pK/BCMVN parameter was taken as the maximum in the values searched per sample. pN = 0.25 for all samples. Out of 60,217 cells, 2,046 (3.4%) were classified as doublets.

### Metacell analysis and filtering

Metacell^41,42^ analysis was performed as described with the following parameter tuning. Feature genes were selected using the functions mcell_gset_filter_varmean and mcell_gset_filter_cov with default parameters. The feature gene set was split into 96 clusters, and clusters were removed if they contained the “undesired variance” genes {*‘Isg15’, ‘Wars’, ‘Ifit1’, ‘Mki67’, ‘Pcna’, ‘Hist1h’, ‘Smc4’, ‘Mcm3’, ‘Top2a’, ‘Fos’, ‘Hsp90ab1’, ‘Hspa1a’, ‘Hif1a’, ‘Xist’, ‘Tsix’*} or had more than 10 genes in the top 100 feature genes correlated with at least two of the UV genes. UV genes and all ribosomal genes were removed from the feature gene set.

Metacells were then constructed using default parameters. Metacells were selected for removal based on a gene footprint score exceeding 1.5 for any of the non-neural/non-cortical markers {*‘Reln’, ‘Lhx5’,’Gad2’, ‘Sst’, ‘Lhx6’, ‘Nrxn3’, ‘Gsx2’, ‘Dlx2’, ‘Aif1’, ‘C1qb’, ‘Hexb’, ‘Igfbp7’*}. All single cells belonging to such metacells were removed from subsequent analyses. Following filtering non-cortical/non-neural cells, metacells were again constructed as above.

### Metacell UMAP and noise cleanup

We removed cell cycle-correlated genes (see also NSC gene module analysis below and Figure S2) and constructed a metacell-metacell similarity graph with the mcell_mgraph_logistic function with mcell_mgraph_max_confu_deg = 7 and otherwise default parameters. UMAP was constructed using 10 neighbors, spread = 1 and min_dist = 0.5. We used MCNoise (https://github.com/tanaylab/mcnoise) to remove background gene expression on the RNA metacell manifold. The parameters used were: num_of_genes_cluster = 13, number_of_mc_clusters = 8, genes_min_diff = 4, thr_max_value = -2, min_number_of_gmctypes_per_batch = 10, min_number_of_batches_per_gmctypes = 5 and excluded 3 gene clusters (2,4,13) from the analysis. The ambient noise levels of all the batches were stable for threshold = -3, and those estimations were used for downstream analysis.

### Inference of metacell temporal flows

Flows were generated using the metacell.flow package^43,56^. This involved inference of optimal flow values given estimated densities of metacells per time point, estimated proliferation rate per metacell and evaluation of differentiation costs (or distances) over the manifold graph.

i. Metacell frequencies per day were estimated directly from the data, by tabulating the time label of all cells and summing across metacells in each day.
ii. Cell-cycle scores (fraction of non-proliferating cells) for metacells were calculated using thresholds of 0.0025 M-gene fraction and 0.001 S-gene fraction^44^. An estimate for the proliferation rate, or the fold-change in number of cells per metacell per time step, was calculated while taking into account two factors: the cell-cycle score, or at what fraction of the maximal rate each metacell is cycling, and an “IPC score”, proportional to the concentration of the *Eomes* mRNA, since it is known that IPCs divide at a slower rate relative to NSCs^46,57^. The formula for the proliferation rate is:

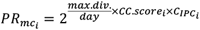

Where the per-metacell cell cycle score 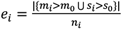 is the proportion of cells in each metacell surpassing the thresholds for M- and S-phase UMI fractions, and the IPC score is 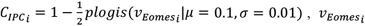 is the *Eomes expression (metacell* e_gc vector) rescaled between 0 and 1 and 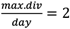, corresponding to a 12hr cell-cycle duration for NSCs.
iii. The construction of metacell differentiation costs were done using the metacell graph described above (used also for UMAP construction).

Given the estimation of i, ii and iii, we built a network model using the function mcell_mctnet_from_mgraph with parameters off_capacity_cost2=10^4^, max_flow_tolerance=0.04, flow_tolerance=0.04, capacity_var_factor=0.2 (and otherwise default). We then solved the flow problems using the function mcell_new_mctnetflow.

### Metacell cell type annotation

Metacells were annotated for cell types based on markers known from the literature. Annotation for intermediate states, namely various classes of immature neurons, utilized marker genes and observed flows from the temporal flow model. We note that metacell construction and temporal flow model inference are agnostic to annotation.

### NSC gene module analysis (Fig 2)

We selected all genes that have max expression (log2 of RNA frequency) of at least -14, and a log2 fold-change of at least 2 between the 5th and 95th quantiles in NSC metacells. The Spearman correlation matrix of these genes’ expression across NSCs was hierarchically clustered and the dendrogram was cut with k=8.

The IPC module was composed of genes with a fold change of at least 2 between mean expression in IPCs and mean expression in NSCs, astrocytes and oligodendrocytes.

The astrocyte gene module was composed of genes with a fold change of at least 2 between mean expression in astrocytes and mean expression in NSCs, IPCs and oligodendrocytes.

The stem gene module was composed of genes with a fold change of at least 1 between mean expression in NSCs and mean expression in astrocytes and IPCs, and had a correlation smaller than 0.5 between their expression across NSC and IPC_cyc metacells and the expression of the cell cycle genes Top2a, Mki67, Mcm4 and Pcna.

The NSC gene module was composed of genes with a fold change of at least 1 between mean expression in NSCs and mean expression in IPC_cycs and astrocytes.

### Single-cell cell cycle analysis (Figs. 2CDE, S2CDEF)

The scRNA matrix was downsampled to 3k UMIs/cell. Single-cell module scores for NSC gene modules 1,3,6,7 and IPC, astrocyte and stem gene modules were the sum of UMIs from each module. We centered the matrix of single-cell cell cycle gene modules and calculated principal components for it. The first two principal components were retained as the cell cycle coordinates, and we rotated them by 140 degrees for convenience. For downstream analyses we assigned as ‘cycling’, those single cells that had total UMIs in cell cycle modules exceeding the 80^th^ percentile. The radial cell cycle phase was calculated for cycling cells as: 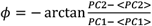 where diagonal brackets denote mean. We used this radial phase to calculate the smoothed phase, using the *principal_curve* function in R with *periodic_lowess* smoothing, on the rotated PC1/2, with cells ordered by the radial phase. Principal curve coordinates were smoothed in a moving average window of 50 points. The phase assigned to each cell was the nearest principal curve point (Euclidean distance). The vector of cells’ phases was binned into 12 equally-spaced bins. Bins 1,4 and 12 corresponded to G1 stage, 5-7 to S, 8-9 to G2, 10-11 to M and 2-3 to G0. While the G0 label was assigned due to a lack of cell cycle signature, we do not claim that every cell with this label is indefinitely post-mitotic, particularly those in progenitor cell types.

### Single cell ordering across differentiation axes (Fig 2E)

To compute the IPC differentiation axis, a principal curve was calculated for the NSC and IPC gene module scores for all single NSCs, IPCs and IPC_cycs. Cells with over 30 UMIs in the astrocyte module were removed from downstream analysis. The developmental axis coordinate for each cell was the principal curve point that was closest to it (Euclidean distance). Cells were then assigned to 20 equally-spaced bins across the axis.

To compute the Astrocyte differentiation axis, a principal curve was calculated for the NSC and astrocyte gene module scores for all single NSCs and astrocytes. Cells with over 30 UMIs in the IPC module were removed from downstream analysis. The developmental axis coordinate for each cell was the principal curve point that was closest to it (Euclidean distance). Cells were then assigned to 20 equally-spaced bins across the axis.

### Low-level filtering of scATAC profiles

Matrices summing ATAC UMIs in single cells in promoter regions ([-2000,200] bp from TSS) and gene bodies (−2kbp from TSS and along gene body) and 500bp around all peaks were constructed. Only scATAC profiles with total UMI counts above the 10^th^ and under the 98^th^ percentiles in all matrices were retained. This removed 14,389/42,141 (34%) of scATAC profiles. In addition, we filtered doublets using AMULET^58^ on each batch separately with default parameters.

### Filtering of non-cortical scATAC profiles

We downsampled the single-cell gene body accessibility matrix *B*_*gm*_ to the minimal library size (7,603 UMIs). We then selected a multi-modal feature gene set that is the union of RNA feature genes and genes variable in their gene body accessibility. We calculated the correlation between the deviation from the mean 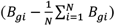 and the normalized expression of the multi-modal feature gene set across RNA metacells (containing non-cortical states, i.e. as in “Metacell analysis and filtering” above). The gene body matrix was clustered based on this correlation, using K-means with K=185, to generate ATAC microclusters. We calculated the correlation between mean microcluster accessibility and RNA metacell expression, and assigned as non-cortical ATAC microclusters those whose top 5 correlated RNA metacells were designated as non-cortical in section “Metacell analysis and filtering” above, and removed all 1,563 (5.6% of total) scATAC profiles belonging to them.

### TSS Enrichment score (Fig S3C)

Only TSSs of genes expressed in the RNA metacell manifold were considered. The ‘flank’ coverage was calculated as the mean marginal UMI count between 900 and 1000 bp (in each direction) from the TSSs. The TSS enrichment score was the fold change of mean coverage in 1bp resolution in 900bp around the TSS, in each direction, over the flank coverage.

### Peak calling

UMIs were pooled from all cells and averaged in 20bp bins. Contiguous genomic intervals with coverage exceeding 195 UMIs, corresponding to approximately 99^th^ percentile coverage, were identified. The intervals were centered on the position with maximal coverage, and uniform-sized peaks were generated by extending from the maximal point and 140bp on each side. Overlapping peaks were unified and re-centered until convergence. This yielded a peak set containing 122,400 peaks, used for all subsequent analyses. We note that peak sets with higher sensitivity were considered but not used, aiming at a conservative high specificity set of epigenetic hotspots for downstream analysis. We classified peaks as TSSs if their distance to an annotated transcription start site was smaller than 1kbp. All other peaks were denoted as CREs.

### Generation of ATAC microclusters (ATAC-MCs)

After filtering low-quality and non-cortical cells, we downsampled the matrix of promoter accessibility to the minimum library size of 2,023 UMIs. For each time point, we clustered the cells into 30 clusters using K-means, based on the correlations between promoter accessibility and fold-change of gene expression from the median across metacells on 1,475 filtered feature genes (essentially identical to those used to generate RNA metacells).

### Matching scATAC profiles to RNA metacells (Fig S3DE)

We defined a linear problem that encodes an optimal assignment of scATAC profiles to RNA metacells. scATAC profiles from time point *t* can only be assigned to RNA metacells containing at least one cell from *t*, since cells for the scRNA-seq and scATAC-seq were sampled isogenically. We calculated the correlation between ATAC-MCs and the fold-change of gene expression from the median across RNA metacells for feature genes (as defined above).

We then constructed a bipartite graph connecting each ATAC-MC with 10 top-correlated RNA metacells, and similarly each RNA metacell with 10 top-correlated ATAC-MCs. This results in an unbalanced graph (with some ATAC-MCs and RNA MCs becoming high-degree hubs). We performed sampling of 10 edges for each MC and each ATAC-MC and combined them to improve balancing. This defines a set of edges (*i, j*) ∈ *E* where i is an ATAC-MC and j an RNA MC.

We can now formulate the ATAC-MC to MC assignment problem linearly, searching for assignments values for each edge *x*_*ij*_ and potential additional over/underflow 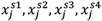. We solve:

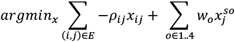

Where *ρ*_*ij*_ is the Spearman correlation linking the accessibility and RNA profiles of the respective ATAC-MCs and RNA MCs, and the over/underflow weights *w*_0_ are parameters (here *w*_1_ *=* −10, *w*_2_ *=* −1, *w*_3_ *=* 1, *w*_4_ *=* 10) approximating a convex cost for imperfect assignment as described below.

This optimization is subject to the following constraints:

i. ∀*ij, x*_*ij*_ ≥ 0 (assigned frequencies are non-negative)
ii. ∑_*i,j*_ *x*_*ij*_ *=* 1 (the total of assigned frequencies is forced to be 1)
iii. ∀*i*, 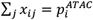 where 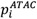 is the observed ATAC-MC frequency
iv. ∀*j*, 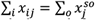 (the total assigned frequency to each MC equals the total ‘frequency capacity’ of each metacell).
v. 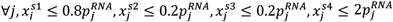where 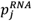 is the metacell frequency for the respective time point. These constraints define ranges of assigned frequency for each metacell, and are matched with over/underflow weights (*w*_0_, see above) such that when 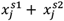 are maximized (negative) cost is minimize. Together with constraint (iv) this encourages the total assignment to an RNA metacell to equal 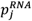.

We solve the linear problem with the ‘lpsymphony’ R package for each embryonic day separately, with the solution giving assigned frequencies from each ATAC-MC to each RNA MC.

### Resampling ATAC metacells given RNA assignments

Given *x*_*ij*_, all cells belonging to ATAC-MC *i* are randomly assigned to metacells according to the proportions *x*_*ij*_/ ∑_*j*_ *x*_*ij*_. This gives an assignment of each valid scATAC profile in the data to an RNA MC. An scATAC matrix for all cells was constructed from the fragments.tsv output of the CellRanger ARC program, using the mcATAC package (https://github.com/tanaylab/mcatac) functions write_scc_reads_from_fragments_file (to create an ScCounts object) and scc_extract (to convert ScCounts->scATAC matrix). Cells were removed if they had less than 6,000 or more than 60,000 UMIs. We then aggregated scATAC profiles according to the RNA MC assignments, using the mcATAC functions scc_project_on_mc and mcc_to_mcatac.

### CRE group annotation (Fig 3AB)

We separated the peak set into TSSs (<1kbp from a TSS of a gene expressed in the RNA manifold) and CREs (all other peaks). Both classes were clustered using k-means (TSSs: K = 10;CREs: K = 60). A subset of CRE clusters was identified to have substantially lower variability across the ATAC manifold, along with qualitatively different sequence content relative to CRE clusters outside of the subset. CREs belonging to clusters in this subset are denoted ‘constitutive CREs’.

### Calculation of putative transcription factor binding motif affinity (Fig 3B)

Sequence affinities were calculated for PWMs of interest as previously described ^59^.

### Selection of neuron branch-specific CREs (Fig 3C)

CfuPN- and IPC-specific CREs: Difference between mean CfuPN (CthPN, SCPN) accessibility and CPN (CPN_L5-6,CPN_L2-3) accessibility > 1 AND absolute difference between mean CfuPN accessibility and IPC accessibility < 1

CPN- and IPC-specific CREs: Difference between mean CfuPN accessibility and CPN accessibility < -1 AND absolute difference between mean CPN accessibility and IPC accessibility < 1

Pan-neuronal CREs: absolute difference between mean CfuPN accessibility and CPN accessibility < 1

CfuPN-specific CREs: Difference between mean CfuPN accessibility and CPN accessibility > 1 AND difference between mean CfuPN accessibility and IPC accessibility > 1 AND NOT in the “Pan-neuronal CREs” group

CPN-specific CREs: Difference between mean CfuPN accessibility and CPN accessibility < -1 AND difference between mean CPN accessibility and IPC accessibility > 1 AND NOT in the “Pan-neuronal CREs” group

### Normalization of ATAC signal (Fig 3E-I)

The following normalization was used for comparing trends in Fig 3E-I. For each CRE the (punctured) marginal coverage was calculated in windows of radius 10kbp around the CRE, and accessibility (UMIs/metacell) was normalized by dividing it by the punctured marginal coverage, while setting the minimum to quantile 0.1 of the punctured marginal coverage, and then linearly transforming the data to the range [0,1].

### Kinetics of branch-specific CREs across neuronal trajectories (barplots above Fig 3EF and under Fig 3HI)

Trajectories were defined as all metacells belonging to defined cell types. For CfuPN: NSC, IPC, IPC_cyc, iCPN/CfuPN, iCfuPN, SCPN, CthPN. For CPN: NSC, IPC, IPC_cyc, iCPN_early, iCPN_late, CPN_L5-6, CPN_L2-3. We defined an NSC-specific gene module as all genes with normalized expression 2-fold higher in NSCs relative to IPCs and glial cell types, and a neuron-specific gene module as all genes with mean expression across all mature neuron cell types 2-fold higher than mean expression in IPCs. The NSC score was the summed expression of NSC genes and likewise for the neuron score. Metacells were ordered and binned (n=20) by the difference between the neuron and NSC scores. These order and binning were used to derive kinetics of CfuPN- and CPN-specific CREs and cell type fractions plotted in figure 3D-I.

### Identification of temporally increasing/decreasing CREs in NSCs (Fig 4C-F)

We derived the per-day accessibility profile of NSCs by multiplying the raw metacell accessibility matrix of NSCs (CREs x metacells) by the matrix of their per-day composition (metacells x days). We transformed this matrix to log fractions, by normalizing by total UMIs per day and taking log2 (plus pseudocount of 1e-5) of the normalized matrix. Temporally increasing CREs were those with a E17-E13 delta ATAC >= 1, and temporally decreasing CREs had E17-E13 delta ATAC <= -1, intersected with the list of CREs with robust methylation signal (as defined below).

### Methylation coverage filtering

We counted 5mC reads in all CREs of the ATAC manifold, and discarded those with less than 20 reads/CRE in all time points, retaining ∼99k/122k CREs.

### Identification of cell type-specific CREs for methylation analysis

NSC: mean accessibility (log-fraction) in NSC >= 0.75 than mean in IPC and >= 0.75 than mean in Astrocytes.

IPC: mean accessibility (log-fraction) in IPC >= 0.75 than mean in NSC and >= 0.75 than mean in mature neural cell types (CthPN, SCPN, CPN_L2-3, CPN_L5-6).

Neuron: mean accessibility (log-fraction) in mature neuronal cell types >= 0.75 than mean in IPC

Astrocyte: mean accessibility (log-fraction) in Astrocytes >= 0.75 than mean in NSC and >= 0.75 than mean in Oligodendrocytes.

### Hi-C – insulation score (Fig 5ABC)

Scores in each time point (E13-E17) were calculated in 250kbp windows, with 1kbp step size, as previously described^37^. High-variance region seeds were 1kbp intervals in which the difference between E17 and E13 insulation was larger than 0.25 or smaller than -0.35. Neighboring seeds were merged and resulting regions smaller than 5kbp were discarded. Mean insulation was calculated for each region in each time point, and the negative of the insulation score was used for visualization (such that higher score == higher insulation).

### Neighboring CRE enrichment in differential insulation groups (Fig 5H)

CREs with mean accessibility >= -15.5 in NSC were selected, their distances from high-variance insulation regions were calculated, and the distribution of neighbors in 1kb distance bins from each type of high-variance insulation region (increasing/decreasing across time points) was calculated. We smoothed the distribution in 100kbp windows and divided it by the number of regions in each group to get the distributions plotted in Fig 5H. To check for significant differences in proximal CRE abundance, all neighboring CREs’ distances were binned into 50kbp-sized disjoint bins, and the distribution of normalized number of neighbors in each bin was compared between insulating and deinsulating regions.

### Normalized SHAMAN D scores between temporally activating CREs in NSCs and cell type-specific TSSs (Fig 5I)

TSSs of cell type-specific TSSs were selected similarly to the procedure in selecting gene modules, where NSC-specific genes were selected based on their differential expression relative to astrocytes and IPCs, IPC-specific genes were differentially expressed relative to NSCs and mature neurons, and astrocyte-specific genes were differentially expressed relative to NSCs and OPCs (all fold-changes of 1). Temporally activating CREs in NSCs were chosen at those whose accessibility per time point had a correlation of at least 0.5 with the time point index, and had a fold change of at least 0.75 between minimum and maximum accessibility per time point. Distances were calculated between temporally activating CREs and cell type-specific TSSs and cut at 1Mbp radius. 2D intervals for SHAMAN D score calculation were generated between the CREs and 10kbp-diameter windows around the TSSs.

### Preparation of training data for accessibility prediction model (Fig 6AB)

We used ICEQREAM (https://tanaylab.github.io/iceqream/) to perform sequence regression on the IPC-NSC ATAC difference, with parameters *max_motif_num=16, n_prego_motifs = 4* and *spat_bin_size = 10*. For each CRE, ATAC UMIs from all CREs within the distance (1kbp,50kbp] of that CRE from NSC scATAC profiles were summed. For each CRE, RNA UMIs from all genes within the distance (1kbp,500kbp] of that CRE from NSC scRNA profiles were summed. All non-constituve CREs with sufficient methylation coverage were split into 11 equally-sized groups, and xgboost (R package) was run with cross-validation on 10 of these groups, with the parameters ‘md’ = 3, ‘eta’ = 0.3, ‘nr’ = 250, obj = “reg:squarederror”, em = ‘rmse’. SHAP values were extracted using the option *precontrib=TRUE*. The R^2 shown is the one obtained by averaging over the R^2 in each of the held-out groups.

### Testing ATAC predictive model of general enhancers

ENCODE SCREEN cCREs (https://doi.org/10.1038/s41586-020-2493-4) were downloaded. All proximal and distal enhancers were collected and unfied into fixed length and no overlap by union and centering. The new interval set was merged with the cortex interval set and total ATAC UMIs in NSCs and IPCs were summed for the combined interval set and normalized. All other sequence and epigenomic features (nucleotide and dinucleotide content, methylation and proximal ATAC and RNA UMIs) were calculated exactly the same as for the cortex interval set, and the xgboost regression model was fit with the same hyperparameters and 10-fold cross-validation.

### Stratification of CREs by E-box affinity, number of proximal elements with high E-box/T-box and methylation (Fig 6CDE)

Distances between all CREs were calculated. E-box and T-box affinity (prego-inferred ‘E_box_1’ and ‘T_box_1’ PWMs shown in Fig S6A) was considered ‘high’ when above the 0.8 quantile. The number of neighboring high-affinity E-box/T-box elements per CRE were capped at 5 each, and 7 total. Active CREs were considered those with mean NSC or IPC ATAC > -16, and the number of neighboring active CREs per CRE was capped at 6.

### MPRA signal cleanup and normalization

Only CRSs that were assigned a p-value of less than 0.1 by the MPRAnalyze pipeline in at least one time point were used. Out of these we selected CRSs whose original genomic intervals overlapped with the CRE set of our ATAC manifold, and were above quantile 0.75 in terms of mean MPRA activity in NSCs or IPCs. MPRAnalyze MAD scores for all time points and cell types were centered (mean 0, s.d. = 1). Two libraries with low QC (matching of MPRA to ATAC signal) were removed.

### Generating xgboost model predictions for MPRA library sequences

Sequence affinities for PWMs obtained from the ICEQREAM model were calculated for all filtered CRSs. We assumed a value of 0 methylation and intermediate-level ATAC (mean value of NSC ATAC) for all CREs. Dinucleotide content was recalculated and the rest of the features were imputed as ‘GC content’ = 0.05, ‘prox_ATAC’ = 10, ‘prox_RNA’ = 10 for all CREs. This aimed at emulating the activated state of the CRE sequence.

