## Supplemental Figures for "Neural stem cell epigenomes and fate bias are temporally coordinated during corticogenesis"

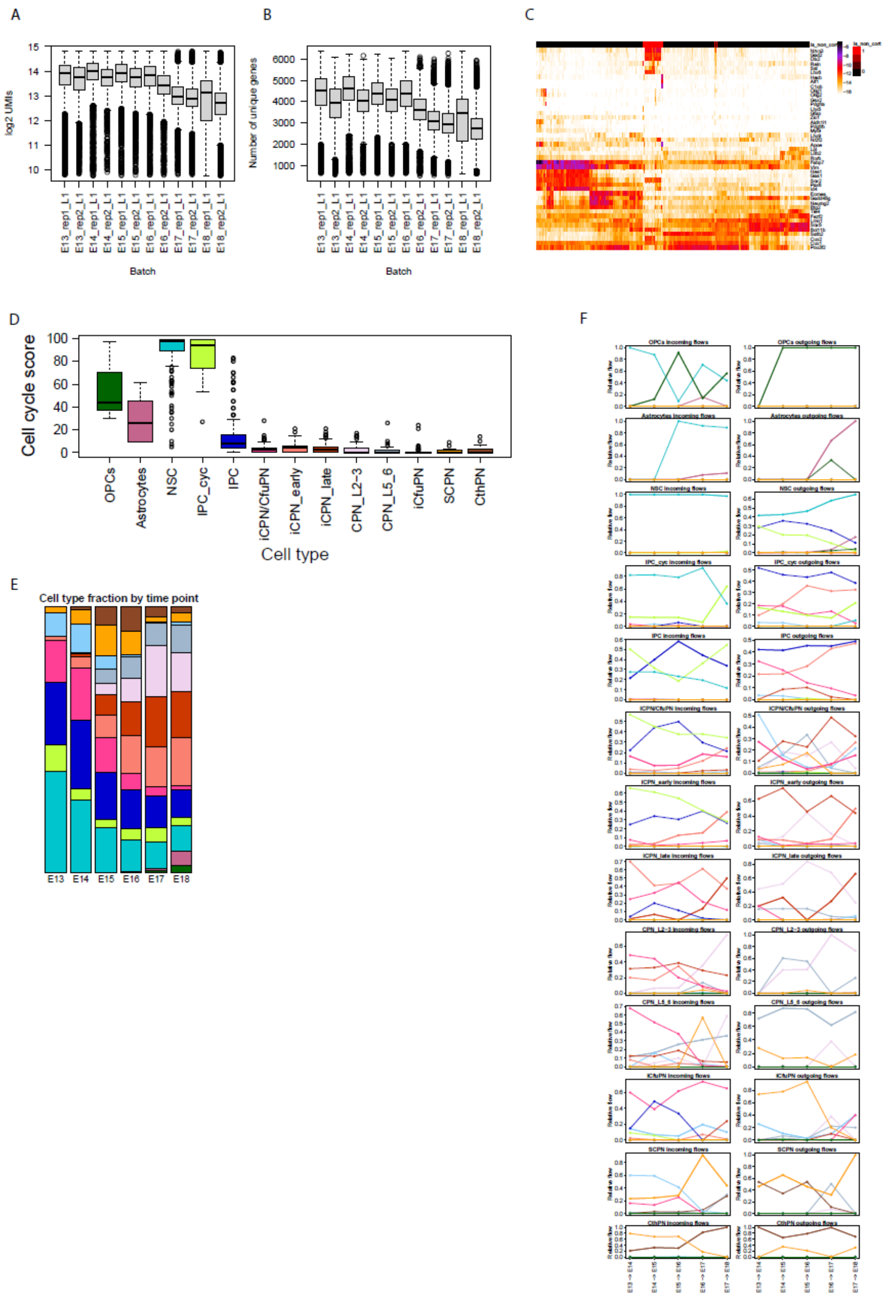


**<u>Figure S1</u>: Additional validation of the scRNA-seq data and the inferred trajectories during mouse cortical development.**

A) Distribution of log2 UMIs in cells per batch in scRNA-seq.

B) Distribution of the number of unique genes per cell in each replicate.

C) Heatmap of expression of non-cortical marker genes in preliminary metacell model to filter out non-cortical cells. Top annotation denotes metacells that were filtered out (red).

D) Distribution of per-metacell cell cycle scores across cell types.

E) Cell type fractions by time point.

F) Relative incoming (left) and outgoing (right) flows aggregated by cell type (line color) by cell type (in subplot titles) across time points. For example, in the fourth row, the main flows going into IPC_cyc metacells in most time points are from NSC metacells, while a minority come from other IPC_cyc metacells, except for the last time point in which NSC and IPC_cyc inflows are almost equal. Outflows are mostly to IPC metacells, and somewhat less to iCPN_early, and finally to iCPN/CfuPN and other IPC_cycs.

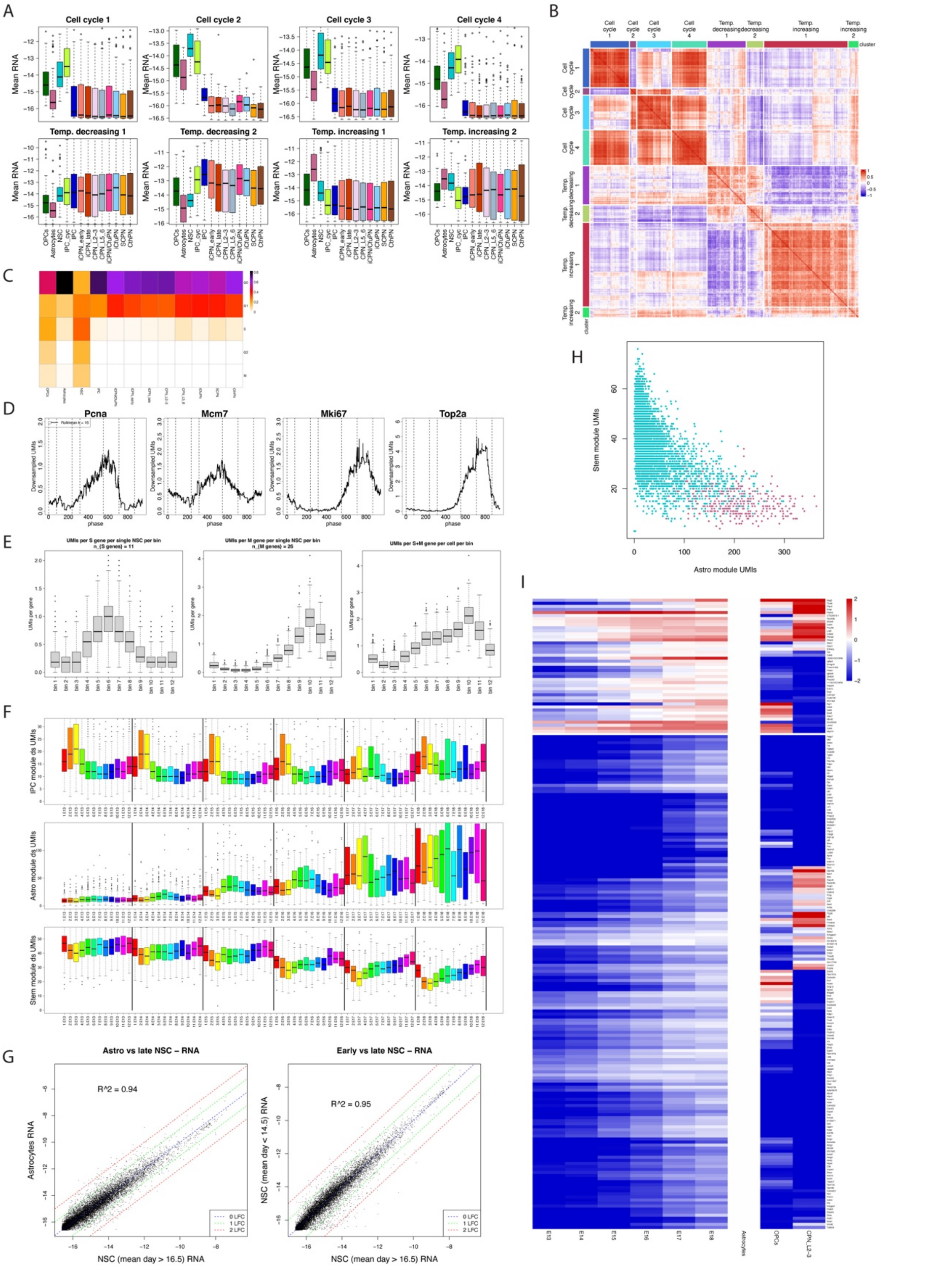

**<u>Figure S2</u>: Correlation between cell cycle and lineages in mouse corticogenesis.**

A) Distribution of expression of genes from each NSC gene module across cell types.

B) Correlation heatmap of NSC gene modules.

C) Cell cycle stages of single cells by cell type. For all cell types except NSC and IPC_cyc, almost all cells are in G0/G1.

D) Smoothed expression (downsampled UMIs) of key cell cycle genes in single NSCs ordered by cell cycle phase.

E) Mean total downsampled UMIs from S-phase and M-phase gene groups (as in Mittnenzweig et al. 2021) across cell cycle bins in NSCs.

F) IPC, astrocyte and stem gene module downsampled UMIs in NSCs, split by time point and cell cycle bin.

G) Comparison of mean expression between astrocytes (left) and late NSC metacells, and between early (right) and late NSC metacells.

H) Stem vs. astrocyte total downsampled UMIs in NSC and astrocyte single cells.

I) Heatmap of gene expression relative to astrocyte levels for all genes temporally activating in NSCs, columns E13-E18 are mean expression in NSCs per day (also relative to astrocyte levels).

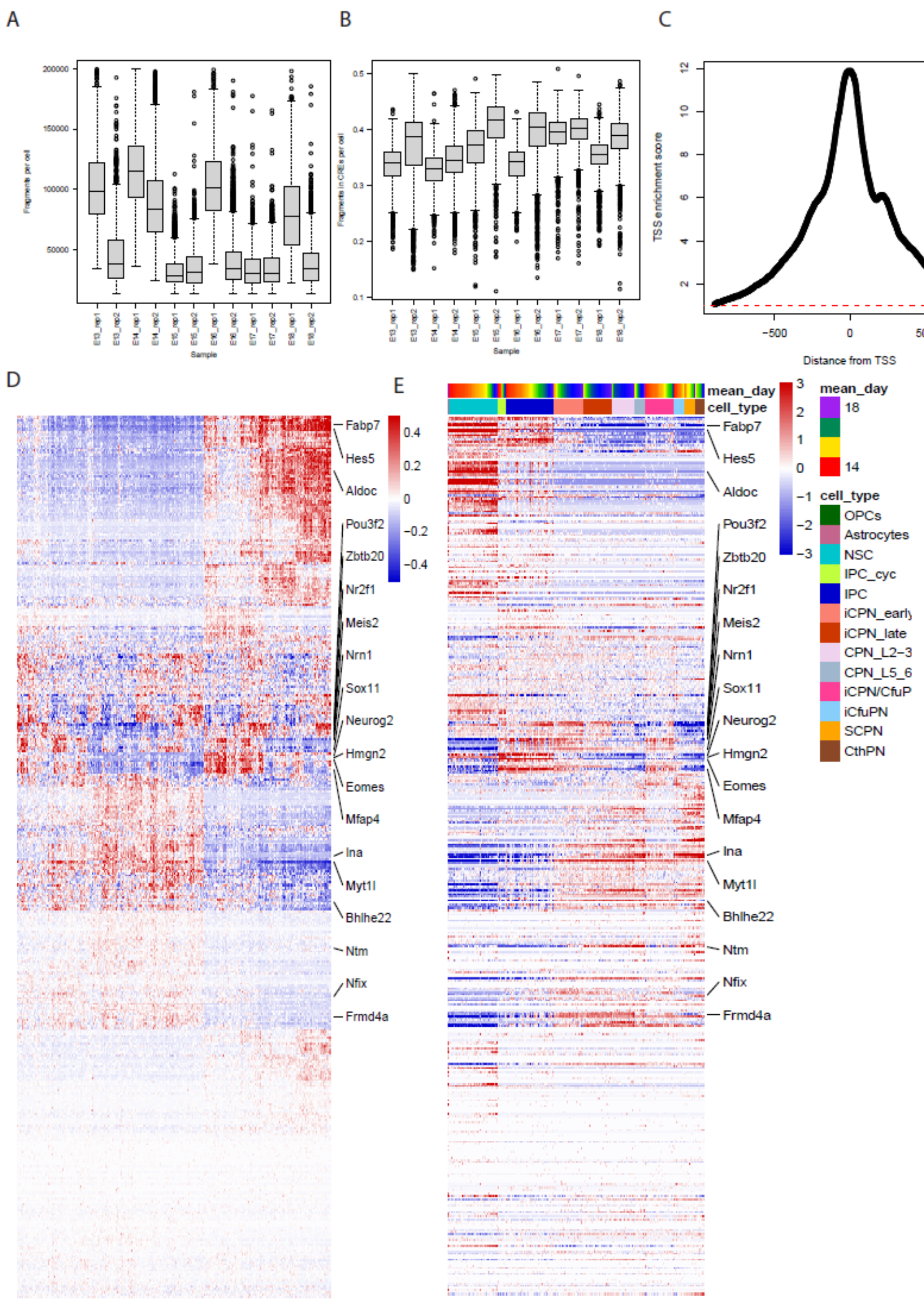


**<u>Figure S3</u>: Additional validation and integration of the scATAC-seq data with the scRNA-seq.**

A) Fragments per cell per scATAC-seq batch.

B) Fraction of fragments per cell in CREs (called peaks) per batch.

C) TSS enrichment score of scATAC fragments (Methods).

D) Relative accessibility of promoter regions of genes used for matching of ATAC and RNA metacells. Marker genes are labeled.

E) Relative expression of genes used for matching of ATAC and RNA metacells.

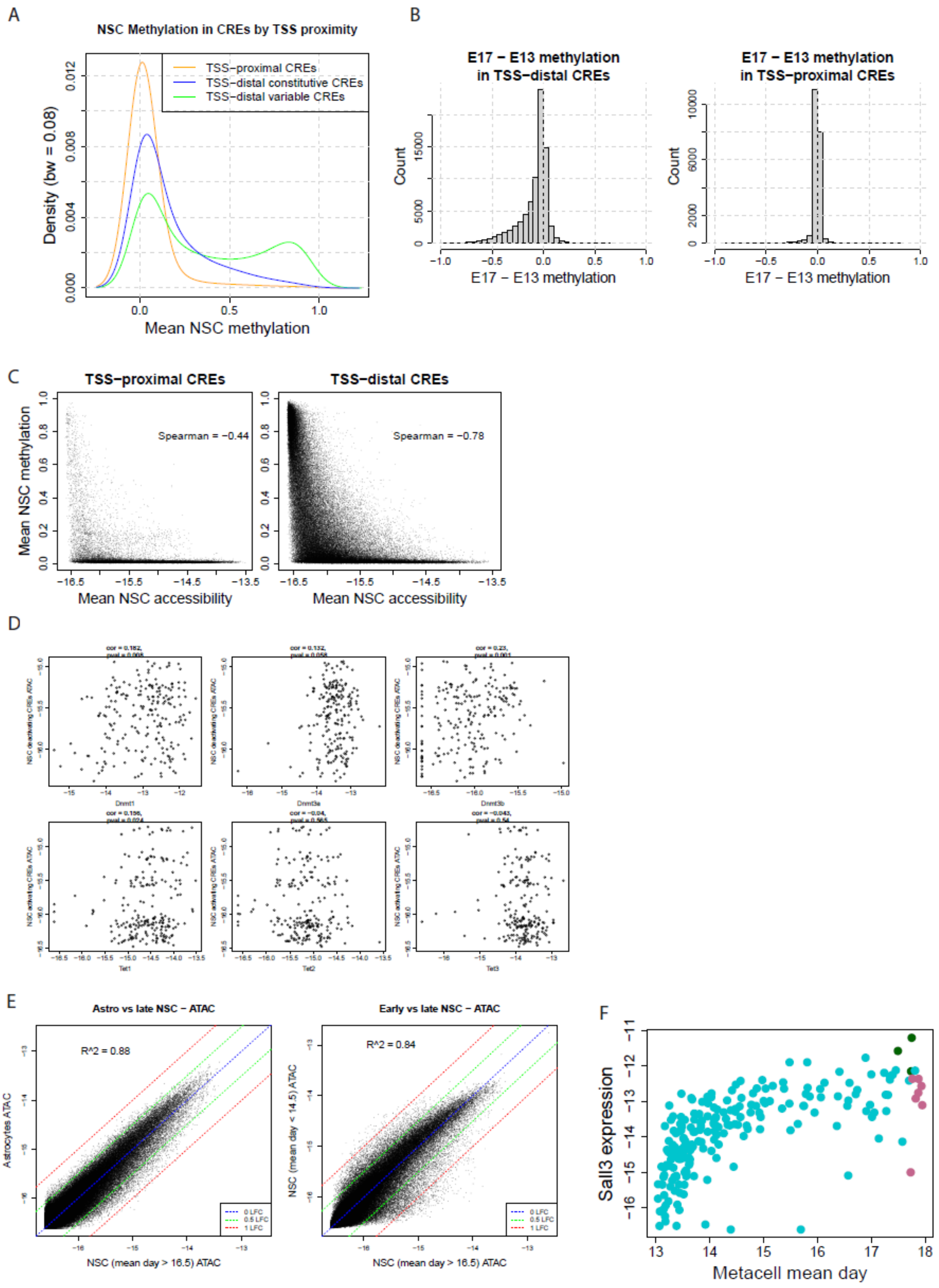


**Figure S4**: **DNA methylation dynamics in NSCs across cortical development.**

A) Probability density of methylation values for different classes of CREs, stratified by TSS proximity and ATAC manifold variability.

B) Distribution of methylation difference between latest (E17) and earliest (E13) time points for TSS-distal (left) and TSS-proximal (right) CREs.

C) Scatter plot of mean NSC methylation vs accessibility for TSS-proximal (left) and TSS_distal (right) CREs.

D) Scatter plots of mean temporally activating CRE ATAC (y-axis) vs. de/methylation enzyme RNA levels (x-axis) in NSCs.
E) Comparison of mean accessibility between astrocytes (left) and late NSC metacells, and between early (right) and late NSC metacells.
F) Sall3 expression by day in NSC, astrocyte and OPC metacells.

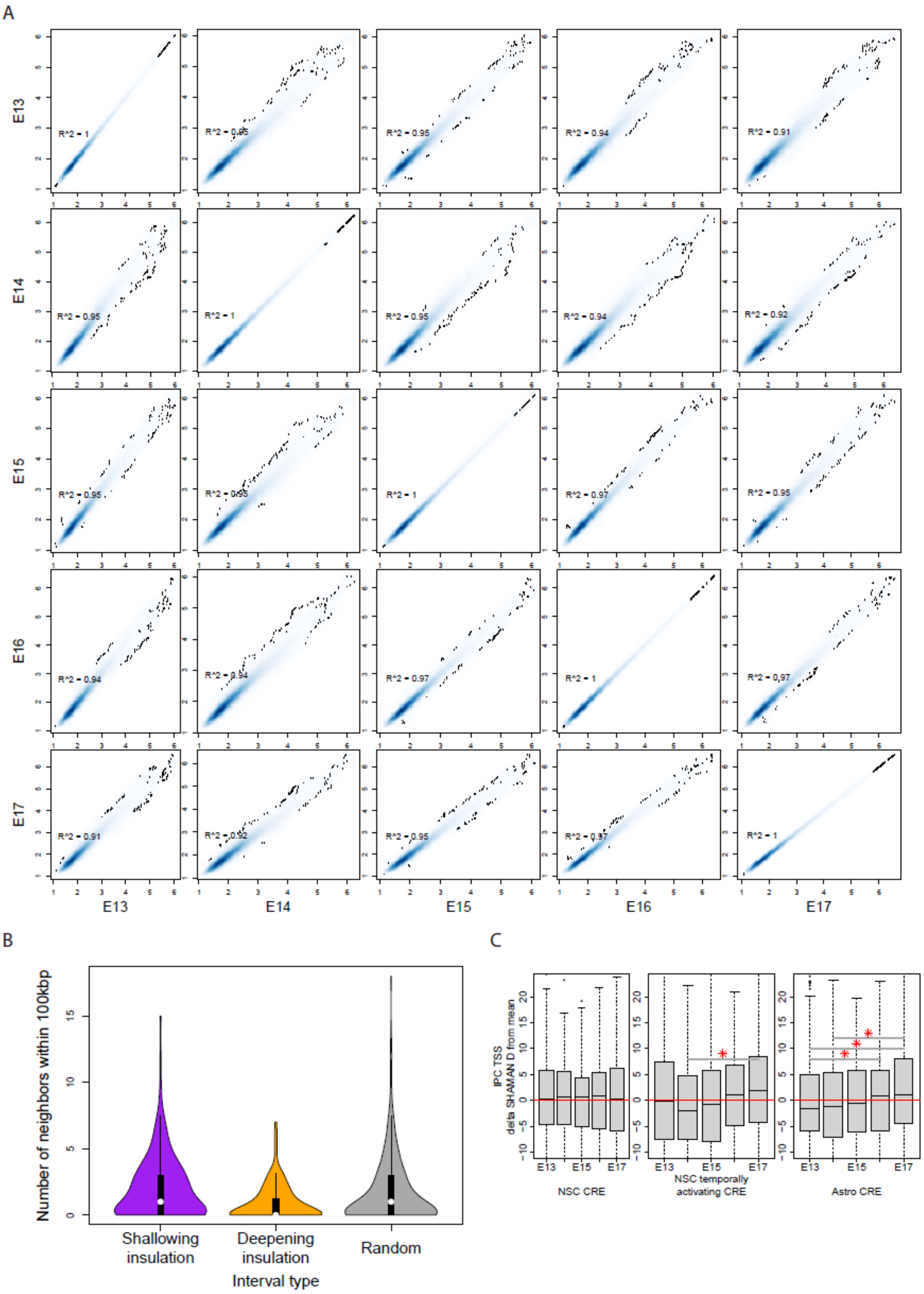


**<u>Figure S5</u>: Chromatin insulation dynamics in NSCs across developmental time.**

A) Correlation of mean insulation scores in 5kbp bins between time points.

B) Number of neighbors within 100kbp of a high-variance insulation region by type of region.

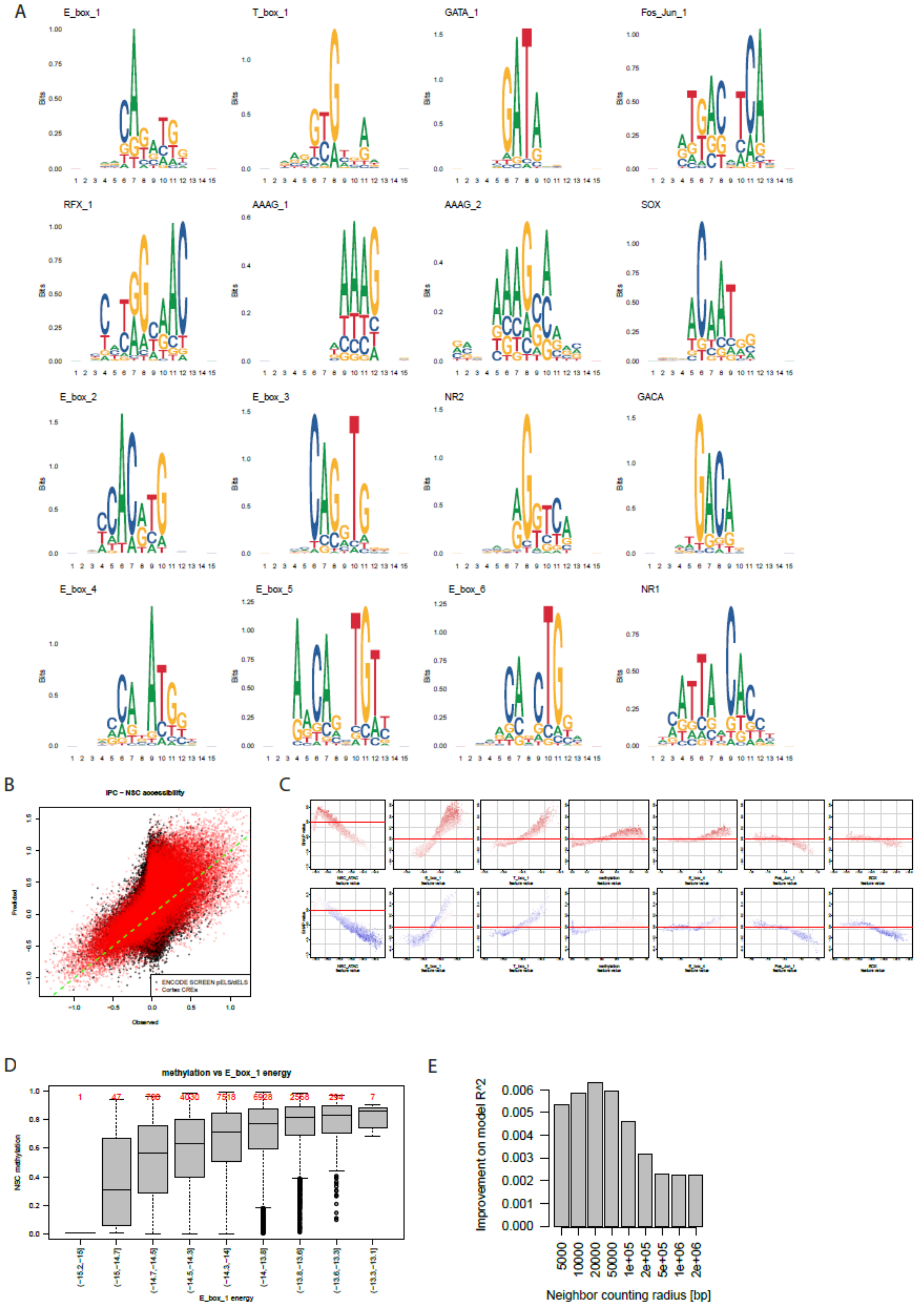


**Figure S6: Inferred TF motifs and correlation to DNA methylation and chromatin accessibility.**

A) Logos of PWMs inferred by ICEQREAM when regressing the mean IPC minus mean NSC ATAC difference (Methods).

B) Scatter of predicted IPC-NSC ATAC for the combined cortex + ENCODE SCREEN interval pool vs. observed (Methods), where cortex CREs are colored in red.

C) SHAP value vs. feature value plots for select features of the xgboost regression model. Points are colored by linearized feature value.
D) Distribution of NSC methylation stratified by E_box_1 motif affinity, showing the two are correlated.
E) Effect on model quality when varying the radius considered for calculating proximal ATAC activity.
